# Predicting the responses of the world’s bird species to increases in agricultural production

**DOI:** 10.1101/2025.02.11.637664

**Authors:** Michela Busana, Ben Phalan, Andrew Balmford, Graeme M. Buchanan, Stuart H. M. Butchart, Alison Eyres, Ashley T. Simkins, David R Williams, Thomas S Ball, Iris Berger, Michael Clark, Lynn V. Dicks, Javier Fajardo, Rhys E. Green, Emma Scott, Paul F. Donald

**Author notes:** Present address: Fondation pour la Recherche sur la Biodiversité, Centre de Synthèse et d’Analyse sur la Biodiversité, Montpellier, France (Michela Busana). **Correspondence**: Michela Busana.

## Abstract

The intensification and expansion of croplands are among the greatest threats to biodiversity, but the likely responses to these changes remain unknown for most species. Using data on responses of 862 bird species to changes in yields of arable crops, we extrapolate vulnerability to agriculture for the world’s other terrestrial birds based on their traits and taxonomy. We estimate that 74-78% of terrestrial bird species globally suffer population declines where natural habitats are replaced by croplands, and that over half cannot persist in cropland at even the lowest current yields. Past impacts of agriculture on birds have been greatest in the tropics, particularly in coastal forest regions of Central and South America and West Africa and in southern and South-East Asia. Using these estimates to model the impacts of future scenarios of agricultural change, we find that continuing current rates of cropland expansion and yield increases (i.e., extrapolating ‘business-as-usual’ trends to 2050) is expected to have more negative impacts on birds (particularly in Central and South America, Eastern Europe, sub-Saharan Africa and South and East Asia) than a potential alternative of a targeted strategy of closing yield gaps while limiting cropland expansion. However, we also identify biodiverse regions of the world where closing yield gaps may have more severe impacts on birds than business-as-usual, such as the Pampas of South America and parts of the Sahel region in West Africa. The strategy of least impact for a particular area can be predicted from the response characteristics of the communities of species present there, offering opportunities to take account of regional context for designing less damaging global food production systems.

## INTRODUCTION

The replacement of natural habitats by agriculture and the intensification of existing agricultural systems are among the most important threats to biodiversity at a global scale (IPBES 2019, Raven & Wagner 2021, Outhwaite *et al*. 2022, Hochkirch *et al*. 2023, IUCN 2024). Globally, a third of agricultural production occurs in areas of high conservation priority (Hoang *et al*. 2023). Agriculture represents one of the most important anthropogenic threats to wild bird populations (Douglas *et al*. 2023, Rigal *et al*. 2023, Fan *et al*. 2024, BirdLife International 2024), and is expected to expand and intensify further in response to the demands of a growing human population and a steadily rising per capita consumption (Duro *et al*. 2020). This will place increasing demand on food-producing regions, which in turn will place huge pressures on biodiversity (Tilman *et al*. 2011, Leclère *et al*. 2020, IUCN 2024). Increasing food system sustainability is critical, and while some gains can be achieved by changes in diet and reductions in per-capita consumption and food loss and waste, a general shift toward more sustainable production, especially in the Global North, will be needed to mitigate the threat posed to biodiversity by agriculture (Foley *et al*. 2011, Scarborough *et al*. 2023).

The projected increasing demand for food can potentially be met by increasing yields in areas already under agricultural production, and/or by increasing the area of productive land, usually at the expense of natural habitats. Real-world experience suggests that both strategies have contributed to rising production in recent decades (although yield increases have contributed far more than expansion; Blomqvist *et al*. 2020), and that both can have significant but often qualitatively different impacts on biodiversity (Tilman *et al*. 2011, Phalan *et al*. 2013, Williams *et al*. 2021).

At present, our ability to predict where agricultural change might generate new hotspots of future extinction risk, or which species might be most threatened, is poor. This is largely because species differ in their responses to habitat loss, land-use change and subsequent intensification of production, with some species being absent from croplands, some being tolerant only of low-yield systems, and others showing positive responses to agricultural expansion or intensification. Recognising that the responses of species to agricultural change may be complex and non-linear, density-yield curve models have been developed to assess species’ relative population densities in natural habitats and on agricultural lands of different yields (Green *et al*. 2005, Phalan *et al*. 2011). Combining these curves with information on arable yields allows assessment of the impact on bird populations of different agricultural production strategies. The broad conclusion of these studies is that, at least for the species assemblages studied, land-sparing strategies are likely to be less detrimental to species overall than land-sharing strategies, provided that (1) a sufficient proportion of the land not converted to agriculture, or restored to natural conditions from agricultural production, is spared for nature (Gilroy *et al*. 2014, Phalan *et al*. 2014, Lamb *et al*. 2016a, Phalan *et al*. 2016, Balmford *et al*. 2019), and (2) that yield increases do not entail unsustainable water and soil depletion or have negative biodiversity impacts beyond the areas where they are applied.

However, constructing density-yield curves is data-hungry and requires dedicated field based surveys to obtain the required information on arable yields and bird abundance, so the responses to different agricultural conversion have thus far been estimated directly for less than 8% of bird species globally (see Results below). It remains unclear whether the observed patterns are representative of bird populations globally, limiting our ability to predict the biodiversity impacts of global scale changes in food production. To address this shortfall, we used an artificial neural network algorithm to extrapolate likely responses for the majority of species for which density-yield curves have not been directly parameterised. This approach classified species into categories of response to agriculture based on known characteristics, such as their morphological and ecological traits. We then combine these extrapolations with models of agricultural expansion to 2050 under two scenarios, business-as-usual and closing yield gaps (Williams et al 2021) to identify areas of the globe where future agricultural production might be most damaging to bird populations. Our aim was to determine where bird communities are most sensitive to the impact of increased agricultural production in 2050, accepting that this is likely to happen despite essential efforts to reduce food waste and promote dietary shifts.

## METHODS

We collated data from studies that previously estimated parameters for density-yield curves for wild bird species, or that generated the data required for us to estimate these parameters ourselves (Phalan 2018). We categorised each observed species into one of three categories based on their likely response to increasing crop production using the characteristics of their density-yield curves. We then derived a generalised density-yield curve for each category and extrapolated these categorisations to all remaining terrestrial bird species by modelling their likely response as a function of their taxonomy and a range of ecological and life-history traits that might be correlated with their tolerance of conversion of natural habitats to agriculture. Finally, we developed a novel set of Area of Habitat (AOH; Brooks *et al*. 2019) maps for terrestrial bird species and combined these with the extrapolated density-yield responses to produce global impact maps to identify areas where birds might be at greatest risk from future crop production under two contrasting scenarios. A graphical representation of our approach is given in Figure S1, and further details of each step in the analysis are given below.

### Deriving density-yield curves

We quantified the relationship between relative population densities of individual bird species and agricultural yields using data from 11 studies across nine countries (Table 1, Figure S2). Taxonomic names and concepts were aligned to match the global bird taxonomy of Handbook of the Birds of the World & BirdLife International (2021). Agricultural yields were quantified from farm surveys primarily conducted on croplands, but also including some data collected from pastures (details on the methodologies are reported in each publication). All studies measured yields in gigajoules per hectare (GJ/ha) except Williams *et al*. (2017), who measured it in kilograms of protein per hectare (kg/ha) (Table 1). In Brazil and Uruguay, the minimum observed yield was 0.1 GJ/ha because all plots surveyed had at least some level of grazing (Dotta *et al*. 2016); plots with an observed yield of 0.1 GJ/ha were treated as comparable with plots in other study areas where the yield was zero.

**Table 1.** Summary of the studies contributing data to the analyses. ‘Years’ indicates the years of data collection, and ‘Species’ indicates the total number of species for which data were sufficient to estimate density-yield curves. The minimum and maximum yields correspond to those observed during the study. The production level is the mean yield required across the whole area to produce the observed level of agricultural production for each study area at the time of the study. ‘Units’ indicates the unit of measurement of yield (note that it differs for Mexico). ‘Natural habitat’ indicates the reference natural habitat(s) against which farmland is compared. Further details of each study are given in the original publications. The last three columns show the proportion of species categorised into each of three density-yield response categories for each study, across all studies for observed species and across all species (n = 9,552) for which categories were modelled. Of the 862 observed species, we randomized the species pool 1000 times to include only once those species observed in multiple studies and report here the mean and standard deviation of the percentages across the randomizations.

| Country | Years | Species | Min.<br>yield | Max.<br>yield | Production<br>level | Units | Natural<br>habitat | Reference | Supersensitive | Sensitive | Resilient |
| --- | --- | --- | --- | --- | --- | --- | --- | --- | --- | --- | --- |
| Brazil & Uruguay | 2010-2012 | 111 | 0.1 | 30 | 2 | GJ/ha | Grassland | Dotta *et al.*<br>(2016) | 0.37 | 0.34 | 0.29 |
| Ghana | 2006-2007 | 167 | 0 | 79 | 19 | GJ/ha | Forest | Phalan *et al.*<br>(2011) | 0.25 | 0.32 | 0.44 |
| India | 2006-2008 | 174 | 0 | 146 | 83 | GJ/ha | Forest | Phalan *et al.*<br>(2011) | 0.27 | 0.18 | 0.55 |
| Kazakhstan | 2009 | 30 | 0 | 23 | 2 | GJ/ha | Grassland | Kamp *et al.*<br>(2015) | 0.30 | 0.50 | 0.20 |
| Mexico | 2011-2014 | 113 | 0 | 4480 | 2000 | kg/ha | Forest | Williams *et al.*<br>(2017) | 0.10 | 0.58 | 0.33 |
| Poland | 2013-2014 | 125 | 0 | 79 | 40 | GJ/ha | Forest & wetland | Feniuk *et al.* (2019) | 0.22 | 0.46 | 0.31 |
| Uganda | 2006-2007 | 256 | 0 | 23 | 10 | GJ/ha | Forest | Hulme *et al.* (2013) | 0.36 | 0.19 | 0.45 |
| UK (Cotswolds) | 2014-2017 | 72 | 0 | 32 | 14 | GJ/ha | Woodland | Finch *et al.* (2020) | 0 | 0.40 | 0.60 |
| UK (Low Weald) | 2014-2017 | 82 | 0 | 22 | 9 | GJ/ha | Woodland | Finch *et al.* (2020) | 0 | 0.44 | 0.56 |
| UK (Salisbury Plain) | 2000-2017 | 75 | 0 | 42 | 8 | GJ/ha | Grassland & woodland | Finch *et al.* (2019) | 0.03 | 0.37 | 0.60 |
| UK (Fens) | 2000-2017 | 97 | 0 | 88 | 46 | GJ/ha | Grassland & wetland | Finch *et al.* (2019) | 0.14 | 0.32 | 0.54 |
| **Overall (observed)** |  |  |  |  |  |  |  |  | **0.28 +/- 0.004** | **0.32 +/- 0.006** | **0.41 +/- 0.004** |
| **Overall (modelled)** |  |  |  |  |  |  |  |  | **0.57** | **0.18** | **0.25** |

To characterise how species’ population densities respond to changes in agricultural yields, density-yield curves were derived for each bird species recorded in each study following the approach of Green *et al*. (2005). Density-yield curves are similar to a generalized linear model for count data with a log transformation of the outcome variable, but they additionally include an exponential parameter, which allows the curve to take concave, convex or bell shapes depending on its value. The explanatory variables included are either the observed yield value or the combination of the yield value and its quadratic term, based on model performance. A formulaic derivation of parameter estimates and the methods used to derive them are given in Appendix S1.

Parameter estimates of each species were extracted for further analyses here. Where parameter estimates were not provided in publications or databases (studies in Brazil, Uruguay and the UK) we re-analysed these datasets using the equations and modelling protocol set out in Appendix S1.

### Categories of density-yield response

We first standardised density-yield curves on the y-axis by dividing each predicted population density value by the mean density in natural habitats (i.e., when yield equals zero). If the predicted density value at zero yields was <0.001/ha, but the maximum predicted density across the range of yields was ≥0.001/ha, we standardised the curve by dividing each density by 0.001. If all predicted densities were <0.001/ha (i.e., if the species was rare in all habitats), we did not standardize the curve to avoid over-inflating the density values that would otherwise tend to infinity. Rare species have similar patterns of changing abundances with yield to common species, but the intercepts of their curves are lower (Phalan et al. 2011).

Next, we followed Green *et al*. (2005) to estimate each species’ relative population density in each study at different yield values and production levels. The method considers the shape of the density-yield response, the predicted densities of the 1,302 species-study combinations across observed yields, and the range of plausible production levels for each study area. The range of observed yields and production levels for each study area are given in Table 1. We defined the minimum permissible yield as the lowest yield capable of meeting the production level if all land within the study area were farmed at that yield. The maximum permissible yield corresponded to the maximum observed yield at each study area. Using the standardized density-yield curve for a species in a given study area, we predicted the relative population density for each species for all yield values between zero and the maximum observed yield. For each study area, we assumed that the area could potentially be covered by different proportions of natural habitats ranging from zero (only agricultural land) to one (only natural habitat). We then calculated the overall population size for each species-study combination at each potential proportion of natural and agricultural land for each plausible production level. For 150 rarely recorded species (those with densities <0.001/ha) we implemented the same steps but used the unstandardized density-yield curve.

Based on the shape of the observed density-yield curves, birds were classified into three categories of response to agriculture: (1) supersensitive species (‘supersensitive’), which were observed only in natural habitats, or in agricultural lands with yields lower than 10% of the maximum yield observed in each study area or below the production level for the study (whichever was smaller); (2) species that persist in agricultural systems but at reduced densities that fall further as yield increases (‘sensitive’); and (3) species that have as high as or higher predicted density in agricultural systems than in natural habitat (‘resilient’) (Figure 1). Further details are provided in Appendix S1. Some species occurred in more than one study, and we categorised these for each study separately to assess species-level variation in response to different types of agriculture and environment.

**Figure 1.**
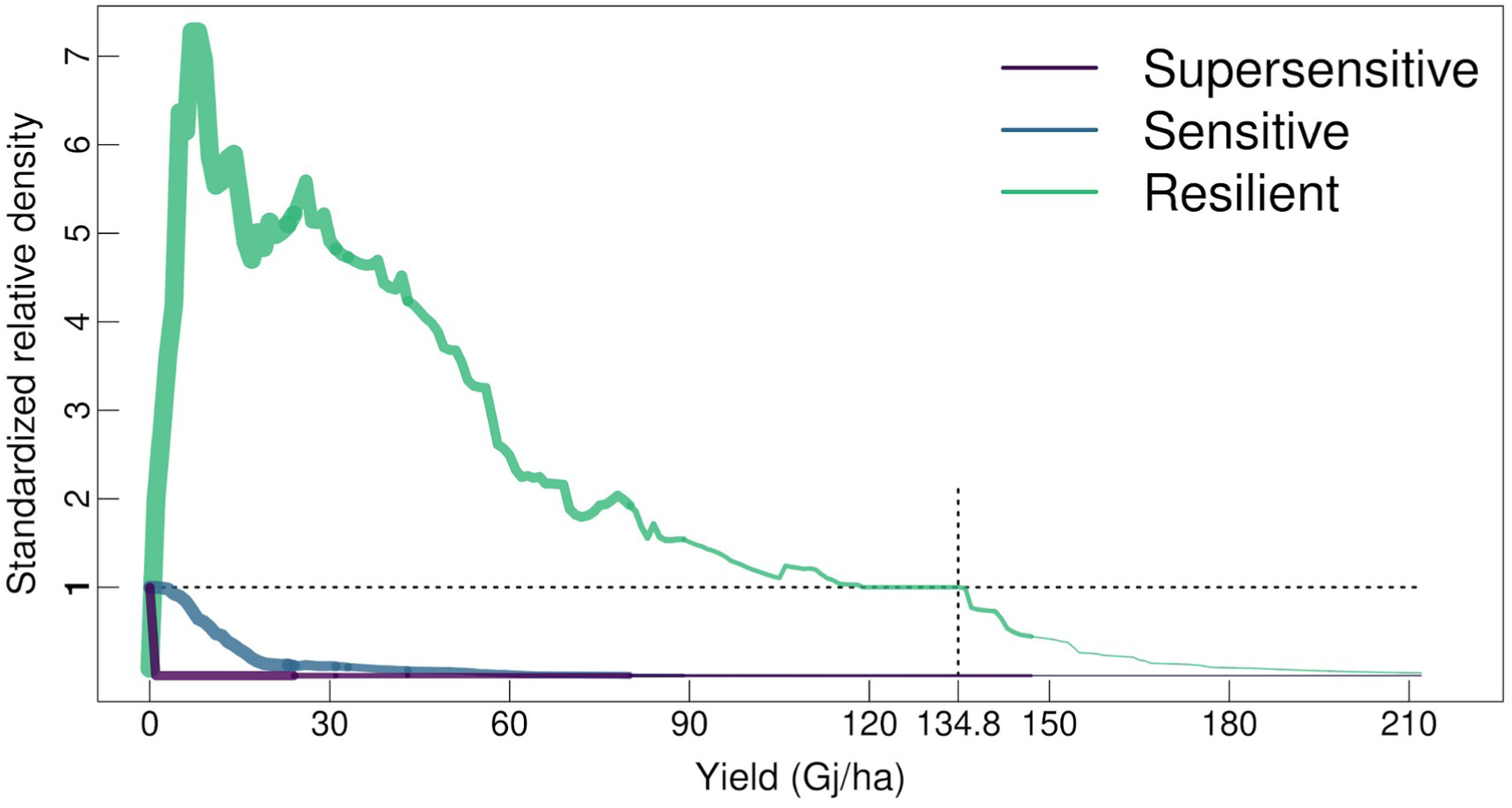
Weighted median species density-yield curves for three categories of response, based on 1,126 density-yield curves estimated from 712 species (we excluded 150 rare species that had predicted densities below 0.001 birds/ha across all yields). Species appearing in multiple studies are down-weighted relative to species appearing only in one. The thickness of the three lines is proportional to the number of species from which it is estimated (see also Figure S5). The maximum observed yield was different in each study, so the number of species informing the line decreases along the *x*-axis. The maximum observed yield in any of the study areas was 146 Gj/ha, and the maximum country-level yield worldwide in 2015 was 134.8 Gj/ha (vertical dotted line), in 2050 under business as usual (BAU_2050_) it was 159.5 Gj/ha, and in 2050 under closing yield gaps (CYG_2050_) it was 211.0 Gj/ha (Williams *et al*. 2021). The relative density is standardized to the observed density of each species in natural habitats (i.e., when yield = 0). Supersensitive and sensitive species had a relative density of 1 in natural habitats, and their relative densities declined as yield increases. Supersensitive species had zero relative densities above 0.1 Gj/ha while sensitive species had close to zero relative densities above 60 Gj/ha. Resilient species had a relative density of 0.08 at zero yields and the highest relative density at yields between 7 and 8 Gj/ha.

Because we derived a response category for each species and each production level in each study area, the response for any species might vary with the production level at a particular area. We assigned a single response category to each species at each study by identifying the most frequently predicted response strategy across the full range of production levels. In the very rare cases where two responses had exactly equal support (four species could have been either sensitive or resilient), we adopted a precautionary approach by assigning the response indicative of a more severe impact of agriculture. Each species-by-study response was analyzed separately to produce an independent density-yield curve and response categorisation. Thus for the 209 species recorded in two or more studies (median = 2, mean = 3.13, max = 9), two or more response categories were derived. To assess the consistency of classification of species across studies we used Kendall’s coefficient of inter-rater reliability for unbalanced datasets after transforming the categories to an ordinal response (1 = supersensitive, 2 = sensitive, 3 = resilient) (Kendall 1962, Landis & Koch 1977).

### Estimating generalised density-yield curves for each response category

Each species in a study is characterized by a unique set of parameter values that define the shape of its density-yield curve, and each study area by a unique value of maximum observed yield. In order to impute generalised curves of changes in relative density with yield for each of the three response categories that can be applied across all species, we grouped the observed density-yield responses across species within each category, excluding rare species that had predicted relative densities below 0.001 birds/ha across all yields, and standardized the curves along the *y*-axis. We then extrapolated the lines up to a yield of 211 GJ/ha, the maximum yield predicted in 2050 by Williams *et al*. (2021). Data from the study in Mexico were excluded as they used a different measure of yield (Table 1). Because extrapolating density-yield curves above their maximum observed yield values might introduce spurious results, we estimated the predicted density value at the highest observed yield (d_obs_) and the expected density value at a yield of 211 GJ/ha (d_max_). If the value of d_max_ was equal to or lower than the value of d_obs_, we allowed the corresponding density-yield curve to predict bird densities above the observed yield. However, when the value of d_max_ was larger than d_obs_, we fixed all the density values above the maximum observed yield at d_obs_.

We calculated the weighted median and interquartile ranges for each set of standardized and extrapolated density-yield responses for each density-yield response category at yield intervals of 0.1 Gj/ha, both at the study level and across all observed species. The weights for each observation were set at 1/f_s_, where f_s_ is frequency with which species *s* was recorded across different studies (range = 1-9).

### Extrapolating density-yield response categories

We used the dataset of 1,302 observed species-study combinations, comprising 862 unique species, to train a multinomial classification model to classify the world’s remaining bird species into one of the three density-yield response categories. We excluded 250 pelagic marine species, 277 species restricted to small islands for which we had no data on agricultural productivity, and 57 species with zero fractional AOH maps (owing to errors in the datasets used to derive AOH maps: see below), leaving 9,552 extant terrestrial species. The model incorporated a range of ecological, morphological and phylogenetic traits that might be correlated with species’ ability to respond to changes in their habitat, including habitat associations, global IUCN Red List category, direction of global population trend, distribution (BirdLife International 2024), ecology, morphology, trophic niche (Tobias *et al*. 2022), life-history (Bird *et al*. 2020) and phylogeny (Jetz *et al*. 2012). Taxonomic order was included as a predictor for all species for which two or more species in the same order were recorded in the observed dataset, with all other species combined into a single ‘rare order’ category. A complete list of the variables included and their sources and definitions is given in Table S1. The complete list of species is available in the archived supplementary data file.

The multinomial classification model used feed-forward deep learning neural networks with 50-fold cross-validation (H2O.ai 2023, R Core Team 2024). We implemented Lasso regularization to avoid overfitting and improve model generalization. Lasso regularization performs feature selection ‘under the hood’ by driving some model coefficients to zero and effectively produces sparse solutions that ignore noisy and irrelevant predictors (Zou & Hastie 2005, Hastie *et al*. 2015). The approach used is robust to multicollinearity, non-normality and missing values in the predictor variables. User-defined model settings for the classification model were tested with cross-validation and are given in Table S2. The model estimates three probabilities for each species, indicating its likely placement in each of the three density-yield response categories; these three probabilities sum to 1 for each species.

### Model evaluation and validation

We evaluated model performance and accuracy using several approaches. First, we extracted commonly used performance metrics derived from the 50-fold cross-validation, including the mean per-class error, log loss, mean squared error, root mean squared error and the confusion matrix highlighting error rates by density-yield response category. Second, we implemented a simplified model with only two density-yield response categories in which supersensitive and sensitive species were grouped together into a single category called ‘disadvantaged’, which was compared against resilient species. The model with only two categories represents a simplified and more parsimonious classification version with the potential to have a higher predictive power. Third, we ran multiple classification models in which the training and test sets were split by country. Species from the same study shared common environmental features that could have influenced their response to agriculture, so it was necessary to assess to what extent it was possible to predict the response of a species in a different study with unknown and potentially different environmental features. The number of species recorded varied greatly between studies, so we combined two or three studies to maintain a ratio of about 80/20 between the sizes of the training and test sets (Table S3).

We then compared the observed or imputed classification of all bird species into density-yield response categories against two independent classifications. First, we compared response categories of birds identified as having high or medium forest-dependence with those of species identified as having low forest dependence or non-forest species (BirdLife International 2024); this classification is not available for other habitat types. Our expectation was that species having medium or high values of forest dependence should be significantly more likely to be categorized as supersensitive or sensitive than as resilient. The second independent dataset used for comparison quantifies bird species’ responses to human pressure on the environment, the Human Tolerance Index (Marjakangas *et al*. 2024), based on the Human Footprint Index (Venter *et al*. 2016). Our expectation was that species classified as resilient to agriculture should have significantly higher tolerance to human disturbance generally than supersensitive or sensitive species.

### Mapping global impacts of agriculture

We mapped global patterns of agricultural production impacts using the three directly observed (for species recorded in the field studies) or imputed (for all other species) density-yield response categories and the generalised density-yield curves for each of those categories, in combination with maps of each species’ global distribution and global patterns of agricultural land and yield (Figure S1). We define impact as the predicted change in birds’ relative density following conversion of unfarmed habitat to agriculture, taking the extent of agricultural area and its yields from the current and future projections of Williams *et al*. (2021). Our models relate only to cropland, since pasture yield values are not available at sufficiently high resolution and because the density-yield curves were parameterised primarily from data collected in cropping systems.

We developed a new set of Area of Habitat (AOH) maps for all the world’s extant bird species at the scale of a grid of 5×5 km pixels. AOH maps describe the spatial distribution of suitable habitat for a species within its geographical distribution (Brooks *et al*. 2019, Lumbierres *et al*. 2022). AOH maps offer a more precise representation of a species’ spatial distribution than broad range maps by significantly reducing errors of commission (Di Marco *et al*. 2017, Dahal *et al*. 2022). We produced AOH maps by combining species’ range maps (BirdLife International & Handbook of the Birds of the World 2021) and data on their habitat preferences and altitudinal ranges (BirdLife International 2024) with a digital elevation model (Robinson *et al*. 2014) and a modified land-cover map of IUCN Level 1 habitat types (Jung *et al*. 2020) at the scale of 100×100 m, using the R package *aoh* (Hanson 2024). Because our baseline was one in which each 5-km pixel is entirely covered in natural land cover, we replaced artificial land cover types (including crops) within the land cover map of IUCN terrestrial habitat classes with the natural habitat classes that would be present in the absence of agriculture (Jung *et al*. 2020; Hengl, Jung, and Visconti 2020).

The AOH maps were re-projected to a grid of 5×5 km (Behrmann projection, ESRI 54017) with pixel values indicating for each species the proportion of apparently occupied area within each pixel. Pixels with a proportional suitability of ≤0.01 were considered to be unoccupied. The breeding and non-breeding AOH distribution of migratory species were combined into a single AOH. Six species among the observed species had a zero fractional AOH (owing to approximation errors or errors in the datasets used to map AOH).

Data on mean crop yields by country (Figure S3) and cropland area (Figure S4) were derived from Williams *et al*. (2021) for the years 2015 and 2050. Two of the scenarios developed by Williams *et al*. (2021) to predict the extent and yields of cropland in 2050 were selected, a business-as-usual scenario (BAU_2050_), which extrapolates recent trends into the future, and a closing-yield-gaps scenario (CYG_2050_), in which yields increase linearly from current yields to 80% of their estimated maximum potential by 2050. Both scenarios yield the same overall level of global food production. The value of 80% was chosen because this approximates the maximum potential in non-trial environments and the economically optimal production level (Williams *et al*. 2021). The CYG_2050_ scenario was predicted to have the lowest impact on terrestrial vertebrates of all the individual scenarios developed by Williams *et al*. (2021), and the one most different from BAU. Because of higher yields, the CYG_2050_ scenario resulted in a lower projected area of agriculture by 2050 than the BAU_2050_ model (20.14 million km^2^ *vs* 27.28 million km^2^) and slightly lower than that in 2015 (21.42 million km^2^) (Figure S4).

For 52 countries that had no available values, crop yields were estimated as the means of their neighbouring countries. No crop yield data were available for 51 small islands or for Greenland or Antarctica (where future crop production was assumed to be zero). The crop yield and agricultural land area (measured as the percentage of land dedicated to agriculture) maps were rasterized to the same 5×5 km grid as the AOH maps and the crop yield in Gj/ha and extracted for each pixel. These yield values were used to calculate the predicted relative density of species in each of the three categories of response to agriculture based on the weighted median of density-yield responses. The crop yield corresponds to the *x*-axis of the density-yield response and the predicted density for each response category to the *y*-axis (Figure 1).

The impact of agricultural change in each 5×5-km pixel was derived as a function of the proportion of the pixel predicted to be covered by agriculture, the yield of the country where the pixels fell, the number of species in each of the three response categories (‘supersensitive’, ‘sensitive’ or ‘resilient’) occurring there and the size of their global distributions. The metric of impact compare the relative densities of species present in each pixel under an assumption of no agriculture (i.e., unfarmed habitats) with those predicted given the agricultural areas and yields of each of the 2015 estimate, and BAU_2050_ and CYG_2050_ scenarios, weighted to account for species-level uncertainty in the allocation of density-yield response categories. The derivation of impact scores is given in algebraic form in Appendix S2.

For each of the three scenarios, we first intersected all species’ AOH maps with the map of crop yields. Each species has three model-estimated probabilities (summing to 1), indicating the likelihood that it is classified as supersensitive, sensitive or resilient, and the generalised density-yield curves for each category predict changes in relative density at different yields. From the intersection of these metrics we extracted the expected change in relative density for each species in each pixel based on its yield. Its value is calculated as the predicted relative density of that species in a pixel covered entirely in natural habitat minus its predicted relative density in the same pixel given the recorded agricultural yield there under that scenario. The latter was derived by taking the modelled probabilities of that species belonging to each of the three density-yield response categories, then weighting the expected relative density of each category at that yield by the estimated probability of the species belonging to that category. Thus, for a species with an estimated probability of being supersensitive of 0.4 and a probability of being sensitive of 0.6, we assigned it the sum of the relative density of a supersensitive and a sensitive species at that yield, weighted by those two probabilities. Each predicted relative density change was weighted by the cover of agricultural land in the pixel in that scenario, the land area of the pixel and the size of the entire AOH distribution of the species (standardised to a range between 1 and 100, since extremely large values of AOH in the denominator reduce the impact on widespread species to near zero). This approach gives higher pixel values to species with small AOH distributions. Positive values indicate that the species is predicted to have a lower relative density in that pixel under the respective scenario than it would in a pixel covered entirely in natural habitat. Negative values indicate that the species could benefit from the presence of agriculture under that scenario. We capped negative values to zero to limit the influence of overall winning species in our calculations, since they do not inform density declines due to agriculture.

For each pixel in each scenario, we summed the relative density changes of all species and calculated the mean impact, its standard deviation and the interquartile range across all species and the percentage of species showing an overall relative density decline in the presence of agriculture in a cell. Overall, we mapped the responses to agriculture of 10,408 species (856 observed from field studies and 9,552 whose density-yield response classes were imputed by modelling).

To identify areas where future agricultural change might have most impact, we subtracted cell values for the 2015 scenario from those of each of the two 2050 scenarios; positive values indicate a larger negative impact on birds between 2015 and 2050, negative values a reduced negative impact. In order to compare the relative impacts of each of the 2050 scenarios, we subtracted the impact scores of the CYG_2050_ scenario from those of the BAU_2050_ scenario; positive values indicate higher impact under BAU_2050_, negative values higher impact under CYG_2050_. Additionally we calculated the proportion of individual species showing a 30% increase and a 30% decrease in impact under those scenarios.

To assess the potential error rate in species classification into categories of density-yield response, we compared the maps of the metrics derived from the three categories of response with those derived from the two-category classification. We separately mapped the impact on threatened species (i.e., species categorised on the global IUCN Red List as Vulnerable, Endangered, or Critically Endangered) to assess their risks. To evaluate the potential influence of AOH size standardisation on the impact of each species, we generated separate maps of overall, mean, and standard deviation impact of birds without applying the AOH standardisation.

## RESULTS

### Bird density changes with crop yields

Density-yield curves were estimated for 1,302 species-study combinations (Table 1), involving 862 unique species. Thus one or more density-yield curves were available for 7.8% of the world’s 10,748 extant terrestrial bird species. On average across all studies, 28 ± 0.4% of species were classified as supersensitive, 32 ± 0.6% as sensitive and 41 ± 0.4% as resilient, although the proportions of species falling into each category varied greatly across studies (Table 1). For the 209 species recorded in two or more studies, and therefore for which two or more categorisations were made, the degree of category concordance was classed as ‘moderate’ (Kendall’s coefficient of concordance, W = 0.54; see Appendix S3 for interpretation and further details). The weighted median density-yield curves for each of the three classes of response are shown in Figure 1 and Figure S5.

### Classification of categories of response

The importance of each of the predictors in classifying species lacking density-yield observations to a density-yield response category is given in Table S4. Cross-validation (80:20) of the model using species for which observations were available suggested that it yielded good discrimination of supersensitive and resilient species but less good discrimination of sensitive species (Table 2). The overall classification success rate was 62%, rising to 68% when supersensitive and sensitive species were modelled as a single category (Table S5 and Table S6 which shows between-country validation results). Modelled density-yield response categories for species not observed in field studies differed in their frequency from observed categories, with 57% of species being classed as supersensitive, 18% as sensitive and 25% as resilient (Table 1). This difference was expected, since observations were biased towards agricultural areas where resilient species were likely to be over-represented, and habitat specialists with small global ranges (which are most likely to be supersensitive) are less likely, on average, to occur in any of the study areas than more widespread species. In models that allocated species lacking observed density-yield response categories to two rather than three classes of response, 80% were classed as ‘disadvantaged’ (i.e., sensitive and supersensitive combined) and 20% as resilient. When we combined observed and modelled species into three or two categories, we estimate that 73.8 and 78.3% respectively of all the world’s birds are disadvantaged by agriculture, with very low or zero densities at even the lowest yields, and 26.2-21.7% are resilient.

**Table 2.** Cross-validation matrix of the model used to impute density-yield response categories for species with no observations, based on 50-folds 80:20 training/test splits of the data for species with observations. Species recorded in two or more studies, which might be assigned to different categories, were treated as independent species. The row total represents the total number of species predicted in that category, while the total number of observed categories can be obtained by summing the values in each row (e.g., 285 observed supersensitive).

|  | **Imputed** |  |  | **Success**<br>**rate** |
| --- | --- | --- | --- | --- |
| **Observed** | Supersensitive | Sensitiv<br>e | Resilient |  |
| Supersensitive | **187** | 52 | 46 | 66% |
| Sensitive | 98 | **205** | 131 | 47% |
| Resilient | 59 | 103 | **421** | 72% |
| Total | 344 | 360 | 589 | 62% |

Our two methods of validation yielded independent support for the observed and modelled density-yield response categories. First, species observed or imputed to be supersensitive or sensitive were significantly more likely to be categorised as having medium or high forest dependence than species categorised as resilient, with 89.6% of medium or high forest dependent species assigned to the supersensitive or sensitive density-yield response categories (Table S8). Second, species observed or modelled to the supersensitive density-yield response category had lower average values of the Human Tolerance Index than species observed or modelled to the resilient category; species assigned to the sensitive category showed a pattern that was somewhat intermediate but more similar to that of resilient species (Appendix S4 and Figure S6). Uncertainties in the classification of observed and predicted species were visualized using a ternary plot (Figure S7).

### Mapping global impacts of agricultural scenarios

Maps of summed species impact by 2015 and in the BAU_2050_ and CYG_2050_ scenarios (all against a baseline scenario of no agriculture) identify the same hotspots of agricultural impact on birds, with the highest impact scores clustering in coastal forest regions of Central and South America and West Africa and in southern and South-East Asia (Figure 2). Mean values of impact, their standard errors and the proportion of species in each cell with impacts greater than zero are shown in Figures S8, S9 and S10 respectively. The interquartile range of impact revealed generally high variability among species inhabiting the same pixels, except in the Sahel and parts of Sub-Saharan Africa, which exhibited the lowest variability between species (Figure S11). Maps of impact based on a two-level category of density-yield response (Figure S12), impact on threatened species (Figure S13), and impact not adjusted by the AOH size of each species (Figure S14) all showed broadly similar geographical patterns.

**Figure 2.**
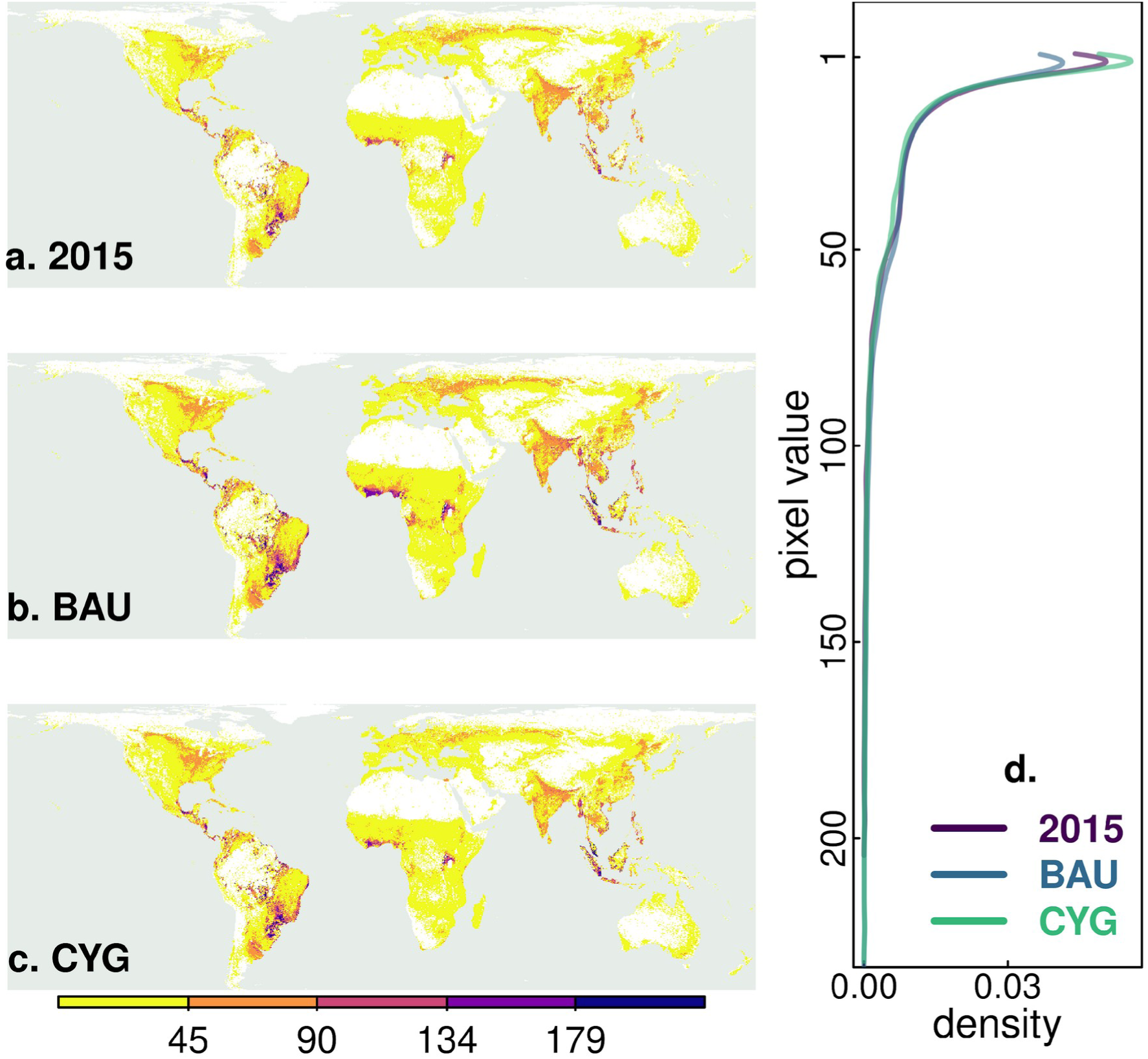
Maps of agricultural impact, summed across species, based on (a) national crop yields and agricultural land area in 2015 and projected yields and agricultural land area in 2050 under (b) a business-as-usual scenario (BAU) and (c) a closing-yield-gaps scenario (CYG). Higher values indicate greater impact of converting natural habitats to agriculture against a baseline of no agriculture. Almost identical patterns are shown when plotting mean, rather than summed, species impact scores (Figure S8). Pixel values were capped at the 0.001 and 0.999 quantile values to reduce the influence of outliers. The graph on the right (d) gives the density of impact scores for each of the three scenarios, showing that CYG has more cells with zero or very low impact compared to 2015 and, particularly, BAU. Histograms displaying the count of pixels by value of each map are shown in Figure S12.

Comparisons between the three scenarios suggest that BAU_2050_ will result in worsening outcomes everywhere compared with 2015, particularly in parts of Central and South America, West Africa and South and South-East Asia (Figure 3a). CYG_2050_ will lead to worsening outcomes compared to 2015 across much of North and South America and western Europe, but improved outcomes across much of Eastern Europe and Central Asia, Africa and South and East Asia (Figure 3b). Summed across species’ entire ranges, 74% of species globally will suffer net negative impact between 2015 and BAU_2050_, compared to 54% between 2015 and CYG_2050_.

**Figure 3.**
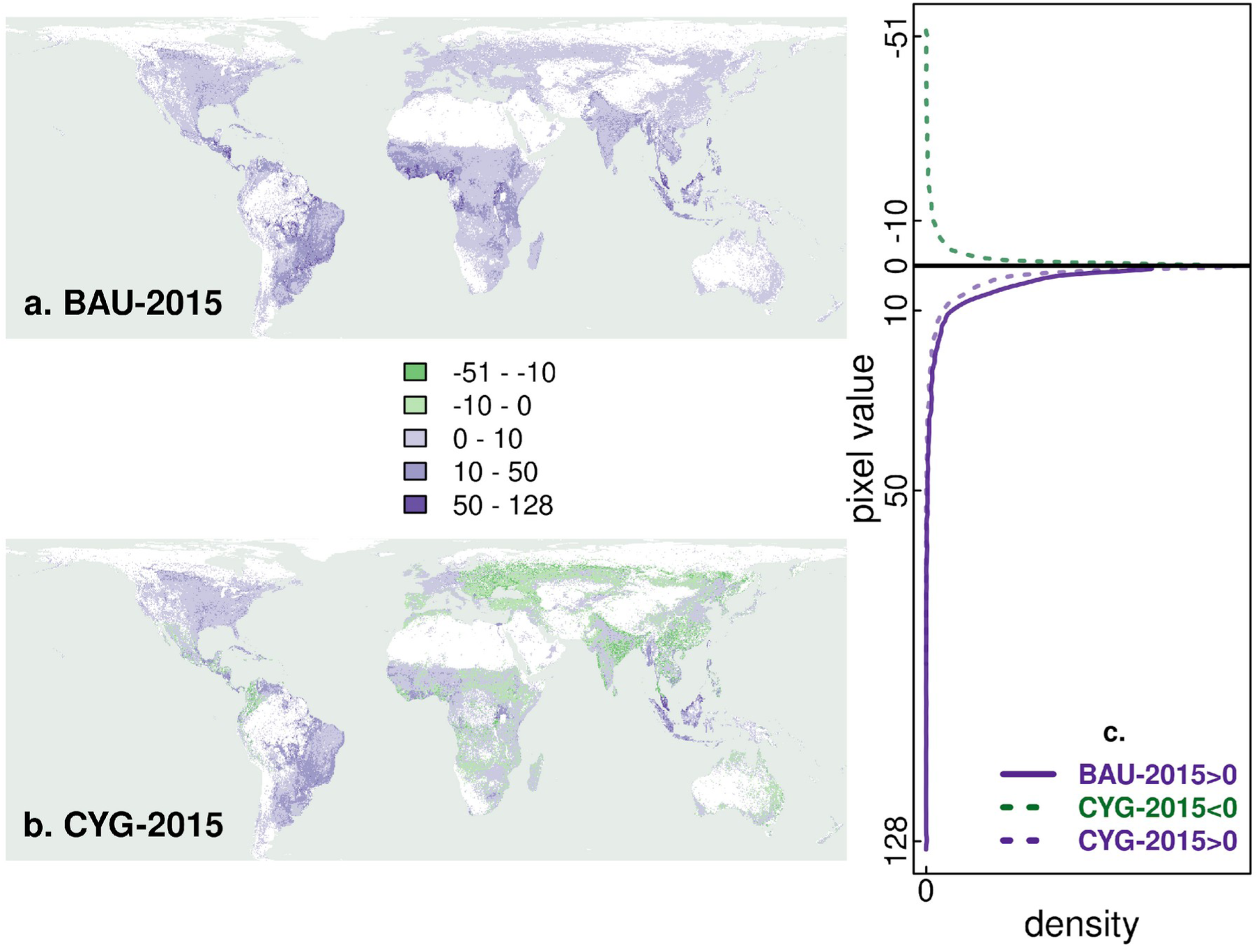
Differences between summed impact scores for (a) BAU and (b) CYG scenarios compared to the 2015 scenario, calculated by subtracting 2015 impact scores from those of BAU or CYG. The resultant differences are binned into five groups. Positive values (purple) indicate an increase in summed impact compared to the 2015 baseline, negative values (green) indicate a decrease. The graph shows a density plot of differences (panel c); note that for the BAU scenario (solid line), all cells had equal or higher summed risk values compared to the 2015 estimate (i.e., there were no negative values). Histograms displaying the count of pixels by value of each map are shown in Figure S13.

When comparing the two 2050 projections, BAU_2050_ was predicted to lead to generally worsening outcomes across much of Central and South America, Eastern Europe, sub-Saharan Africa and South and East Asia, but improved outcomes in parts of southern South America and West Africa (Figure 4a). Differences in impact between BAU_2050_ and CYG_2050_ were generally of small magnitude, so we assessed the spatial distribution of bigger differences between the two scenarios. The proportion of species predicted to suffer a >30% higher impact under BAU_2050_ compared with CYG_2050_ (Figure 4b) was generally higher than the proportion of species predicted to suffer a >30% higher impact under CYG_2050_ compared with BAU_2050_ (Figure 4c).

**Figure 4.**
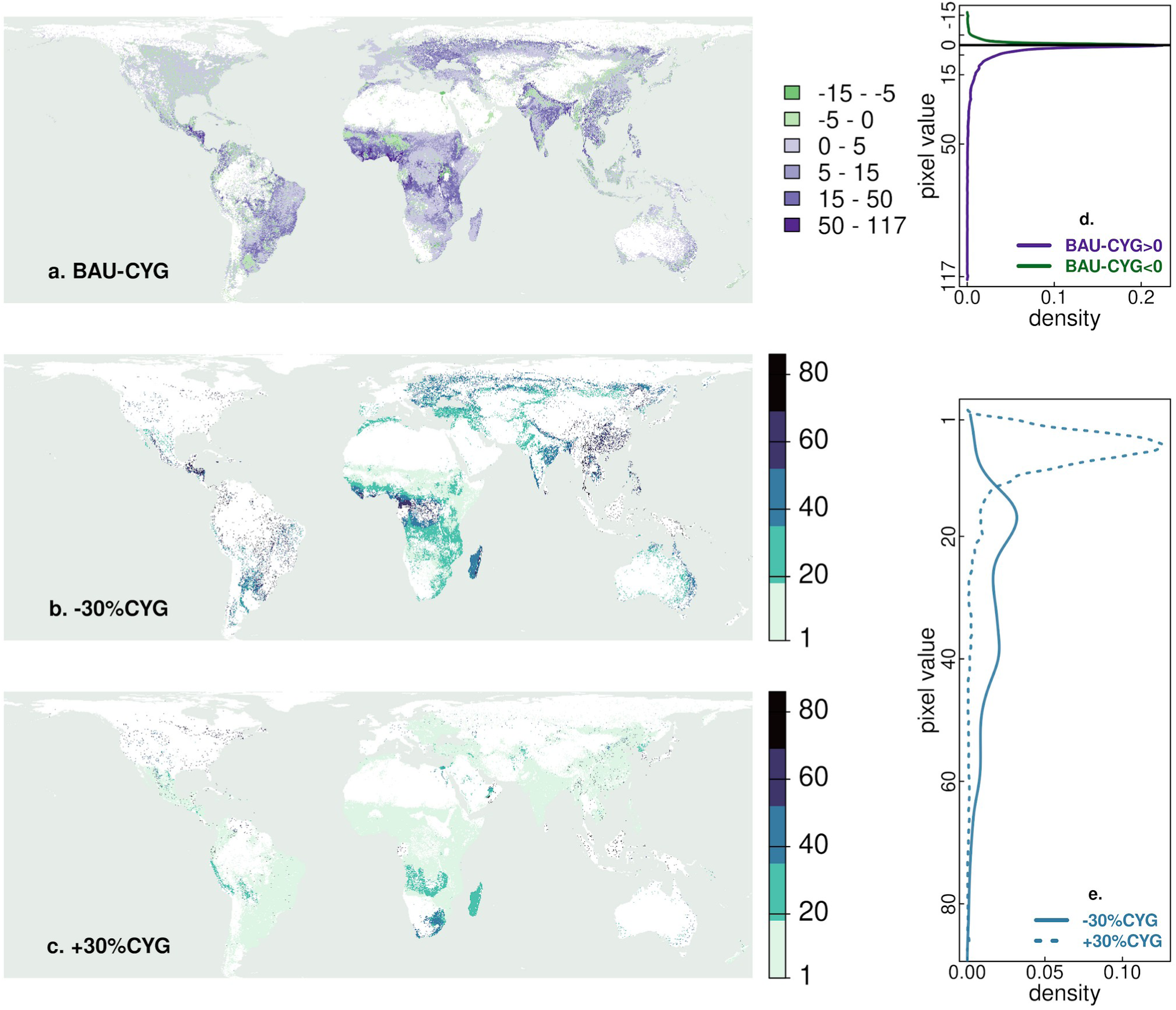
Comparisons between the 2050 business-as-usual (BAU) scenario and the 2050 closing-yield-gaps (CYG) scenario. (a) Differences in summed impact scores, calculated by subtracting the summed CYG impact score from the summed BAU impact score, with positive values (purple) indicating a higher impact under BAU and negative values (green) indicating a higher impact under CYG. (b) The percentage of species predicted to have a >30% lower impact score under CYG compared to BAU. (c) The percentage of species predicted to have a >30% higher impact score under CYG compared to BAU. The graphs on the right show density plots of differences. Histograms displaying the count of pixels by value of each map are shown in Figure S14.

## DISCUSSION

Species traits have been used successfully to model the responses of wild birds to the conversion of natural habitats to agriculture in some locations (e.g., Binley *et al*. 2023), but this analysis is the most comprehensive attempt to do so for the great majority of the world’s terrestrial bird species. We find that up to 78% of the world’s bird species are likely to be sensitive or supersensitive to agriculture, predicted to have low or zero population densities in even the lowest-yielding production systems. The remaining 22% are resilient to some extent, predicted to occur in higher densities in agriculture than in the natural habitats that it replaces, although many of these species are likely to have greatly reduced densities at current yields that are predicted to fall further at the highest yields projected for many countries in 2050. These patterns dictate that most of the world’s agricultural lands support species-poor bird communities at current yields, with further increases in yield having relatively little additional impact, whereas the expansion of new agriculture into natural habitats will reduce or extirpate the populations of the majority of species found there.

Our cell-level metric of global impact suggests that agriculture has had the greatest overall impact on bird populations in parts of Central and South America, tropical West Africa and much of Southern, Eastern and South-Eastern Asia (Figure 2a). The areas of greatest summed impact tend to be tropical areas of high avian endemism (Stattersfield *et al*. 1998), particularly the Brazilian Cerrado and Atlantic Forests in South America, the Isthmus of Panama in Central America, the Lower Guinean Forests of West Africa and the Sundaic Forests of South-East Asia. The same areas are expected to be those most severely further impacted by 2050 under any scenario (Figure 2b,c). A similar pattern of greater impact of agriculture in tropical regions was reported by Zhao *et al*. (2024) and results from the high number of small-range forest specialists in the tropics and the more extensive areas of open, more agriculture-like habitats (temperate grasslands, savanna, tundra, steppe etc.) across much of the drier and colder temperate zone. Another factor that could explain spatial variation in the vulnerability of species to agriculture is the much longer history of extensive agriculture in some regions than others. Large-scale agricultural practices were present across much of Europe before 800 CE, whereas across most of South America croplands only began to emerge after 1700 (Pongratz *et al*. 2008). As a result, species in Europe may have had more time to evolve with and adapt to agricultural landscapes (or be driven to extinction) compared with those in the tropics (Balmford 1996). Furthermore, the ranges of species used to develop the AOH maps might not reflect pre-agricultural distributions in areas where agriculture has been dominant for centuries, with some species’ populations likely to have been extirpated from these areas.

The trajectory taken to meet food requirements by 2050 will significantly influence the extent to which additional environmental damage will occur. Our global approach offers the potential to develop a strategy that targets the closing of yield gaps to landscapes where increasing yields would have the greatest net benefit, while also identifying and appropriately safeguarding agricultural systems where further yield increases might be most damaging to biodiversity. We predict that under a business-as-usual scenario, there will be an additional net negative impact on birds across all the world’s agricultural land (Figure 4a). In contrast, a closing of yield gaps and corresponding reduction in cropland area (Williams *et al*. 2021) is predicted to reduce impacts across extensive parts of Eastern Europe, Central, Eastern and Southern Asia, Australia and Sub-Saharan Africa. Our results therefore provide global-level support for previous suggestions (e.g., Balmford *et al*. 2005, Phalan *et al*. 2016, Tilman *et al*. 2017, Phalan 2018) that future impacts on natural systems might be reduced by a carefully targeted closing of yield gaps, relative to an extrapolation of recent patterns of change (business-as-usual). Such a strategy should not be equated with an endorsement of all high-yield practices; rather it is a recognition that in some parts of the world there is the potential to increase yields above their current rates of increase with relatively little additional impact on bird populations, due to the response characteristics of the species occurring there.

For any benefits to accrue from such a strategy, a sufficient proportion of the land not converted to agriculture, or removed from production, would need to be spared for nature, and higher yields, usually leading to higher profits, should not stimulate additional land conversion and over-production. This pattern may not apply to other taxonomic groups, and our models take no account of the sustainability of yield increases in terms of depletion of soil nutrients and water, carbon balance or of the consequences of filling yield-gaps for surrounding areas, for example through pesticides or run-off of fertiliser nutrients into watercourses (e.g., Ma *et al*. 2021, Penuelas, Coello, and Sardans 2023, Li *et al*. 2024). Nevertheless, our findings that (i) across much of the world’s farmland current levels of yield are already so high that all but the most resilient species have already been extirpated, and (ii) that a high proportion of species in remaining natural habitats are sensitive or supersensitive to agriculture at even the lowest yields, dictate that if agricultural production needs to rise, the priority in most areas is to limit the loss of natural habitats. They also suggest that if the compensatory need for increased yields elsewhere is targeted to areas where their impact will be lowest, significant further loss might be averted.

Closing yield gaps to attainable levels to meet projected demand in 2050 could potentially help spare an area equivalent to that of the Indian subcontinent from conversion (Phalan *et al*. 2014) and could result in greater above-ground carbon storage than clearing land for further production (Gilroy *et al*. 2014, Lamb *et al*. 2016b, Williams *et al*. 2018). It might also be a particularly efficient way to meet rising food demand; for example in non-irrigated systems, average cereal yields typically reach less than 50% of their potential yield (Lobell *et al*. 2009) and closing yield gaps could increase crop calorie production by up to 80% compared with levels in 2000 (Pradhan *et al*. 2015). However, our results also identify areas of the world where a strategy of closing yield gaps might have greater negative impacts than continuing current patterns. These patterns may be influenced by the predominance of resilient species along with a combination of optimal cropland extension and yields that support these species in these areas. Our results therefore support concerns that a singular focus on closing yield gaps may be inappropriate due to complex heterogeneity in production systems and the responses of biodiversity (Cunningham *et al*. 2013, Phalan 2018).

A number of caveats apply to our analyses. Additional field research is needed to gather more data on both intraspecific and interspecific species densities in natural habitats and across a range of crop yields and management systems. This will help validate our classification approach for a broader range of species in diverse habitats and enhance the accuracy of our predictions. Our analysis excludes pastures and does not account for their potential negative effects on birds’ population densities (e.g., Godoi *et al*. 2018). Future studies are needed to incorporate the impacts of both crops and pastures on bird populations at a global scale. Furthermore, our projections assume that natural habitats are intact and capable of supporting sensitive and supersensitive species at their maximum potential densities. This assumption may not hold true, as many unmeasured factors such as climate change, pollution, hunting and other types of land-use changes could adversely affect population densities in natural habitats. Our cell-level metrics of impact do not account for the role of habitat fragmentation, which could exacerbate the effects of farming on bird densities in natural habitats, potentially hindering the ability of species to persist in natural habitats that have not been converted to agriculture (Lamb *et al*. 2016a). There is also a need to extend our approach to non-avian groups, in order to represent biodiversity more generally.

Further work is also needed to assess land-use and spatial planning policies in the countries and regions within them that our results highlight as being particularly sensitive to further conversion of natural habitats, and how best to influence those policies and plans. These will need to consider whether maintaining low yields in one region or country may lead to agricultural expansion elsewhere, thereby ‘offshoring’ the environmental costs of food production (Boakes *et al*. 2024); perhaps over 50% of the biodiversity loss associated with consumption in developed economies occurs outside their territorial boundaries (Wilting *et al*. 2017, Schwarzmueller & Kastner 2022).

Successfully navigating these challenges will allow appropriate land management activities to be targeted towards places where doing so will deliver the greatest benefits for biodiversity. In parallel, further work is needed on how yields can be raised with the least impact on biodiversity. Experience has shown that simple, low-cost measures, such as integrating patches of natural habitats within crops or providing artificial nest sites, can enable some species to persist or even thrive in high-yield food production systems (e.g., Schmidt *et al*. 2017, Lindell *et al*. 2018, Ke *et al*. 2023), effectively manipulating their density-yield curves to increase population density at high yields. Recently developed agroecological systems designed to be high yielding without degrading ecosystems, can also change the shape of density-yield curves (Berger *et al*., in review). A full consideration of the needs of resilient and also sensitive species through ‘sustainable intensification’ should form an integral part of strategy that advocates a closing of yield gaps (Garnett *et al*. 2013, Cassman & Grassini 2020) and would contribute to Target 10 of the Kunming-Montreal Global Biodiversity Framework (‘Enhance Biodiversity and Sustainability in Agriculture, Aquaculture, Fisheries, and Forestry’). At the same time, progress needs to be made towards Target 16 (‘Enable Sustainable Consumption Choices to Reduce Waste and Overconsumption’), which would reduce the need to increase productivity at all.

### Data statement

The data used in the analysis can be downloaded from the following sources:

● land-cover: Jung *et al*. (2020); Hengl, Jung & Visconti 2020;
● digital elevation model: Robinson, Regetz, & Guralnick (2014);
● range maps: Bird species distribution maps of the world. Version 2021.1. Available at http://datazone.birdlife.org/species/requestdis
● IUCN Red List assessments: BirdLife International (2021) IUCN Red List for birds. Available at https://datazone.birdlife.org/species/search
● density-yield curves: see references in Table 1. Parameters of the density-yield curves and median predictions at different yield values are also available on [archive address];
● species informations including taxonomy, inclusion/exclusion from the analyses, density-yield response categories with probabilities: available on [archive address];
● traits: Tobias et al. (2022); Bird et al. (2020); BirdLife International and Handbook of the Birds of the World (2021); BirdLife International (2024);
● raster files: archived on [archive address].

The results were obtained using R (R Core Team 2024) and GDAL/OGR (2024). All software to reproduce the results and Figures is archived on [archive address]. Additional details can be found in Appendix S5.

## Acknowledgements

This work was supported by the CCI Collaborative Fund, which is funded by Arcadia (a charitable fund of Lisbet Rausing and Peter Baldwin), the Rothschild Foundation, the A.G. Leventis Foundation, the Isaac Newton Trust, and the Prince Albert II of Monaco Foundation. We thank the 4C group at the University of Cambridge and M.W. Dales for their support and for providing access to the Sherwood high performance computing cluster. We thank Tom Finch for providing the UK datasets and supporting their analysis. MB received funding from the French Foundation for Biodiversity (project Acoucene) since July 2023.

## Author contributions

All authors: methodology, interpretation, conceptualization, investigation, formal analysis, writing (review and editing). MB, BP, PD: bird data curation. MB, DW, MC: farmland data curation. MB, PD, BP, DW, AB: agricultural scenario investigation. MB, SB, AE, GB, PD, AS: AOH maps preparation. MB: data analysis and software implementations. PD and MB: writing (lead)

