## supplementary online material for "Predicting the responses of the world’s bird species to increases in agricultural production"

#### Contents

**Figure S1: Graphical representation of workflow** **2**

**Figure S2: Study sites location** **3**

**Figure S3: Estimated mean crop yields by country** **4**

**Figure S4: Estimated agricultural land** **6**

**Figure S5: Curves relating species relative standardized density per hectare (ha) to agricultural yield for different response categories** **8**

**Figure S6: Comparison between the density-yield response categories and the tolerance to human pressure** **9**

**Figure S7: Ternary plot** **10**

**Figure S8: Mean impact in the three scenarios** **11**

**Figure S9: Standard deviation of the mean impact** **12**

**Figure S10: Proportion of species with projected density declines** **13**

**Figure S11: Interquartile range of the impact** **14**

**Figure S12: Impact maps based on two density-yield response categories** **15**

**Figure S13: Impact maps based on globally threatened species only** **16**

**Figure S14: Results without the correction with AOH size** **17**

**Figure S15: Histogram of cell values in 2015, BAU and CYG** **19**

**Figure S16: Histogram of cell values in (BAU minus 2015) and (CYG minus 2015)** **20**

**Figure S17: Histogram of cell values in (BAU minus CYG)** **21**

**Table S1: Variables used for modelling density-yield response categories** **23**

**Table S2: User-defined model settings for the classification model** **27**

**Table S3: Country-wise validation of species classification** **28**

**Table S4: The importance of the different predictors in classifying species to density-yield response categories** **29**

**Table S5: Classification into two-categories of response to agriculture** **33**

**Table S6: Country-wise results of species classification into three categories of response to agriculture** **34**

**Table S7: Comparison with forest-dependent species** **36**

**Table S8: Mathematical symbols** **37**

**Table S9: Classification of species observed in multiple sites** **38**

**Appendix S1: Estimating density-yield curves** **39**

Additional details about the classification of species into categories . . . . . . . . . . . . . . . . . . . . 39

**Appendix S2: Formulaic definition of calculation of cell-level metrics of impact** **41**

**Appendix S3: Concordance between multiple assessments of density-yield category of the same species** **43**

**Appendix S4: Validation of density-yield response categories using the Human Tolerance Index (HTI) of Marjakangas et al. 2024** **44**

**Appendix S5: Software details** **45**

**References** **46**

### Figure S1: Graphical representation of workflow

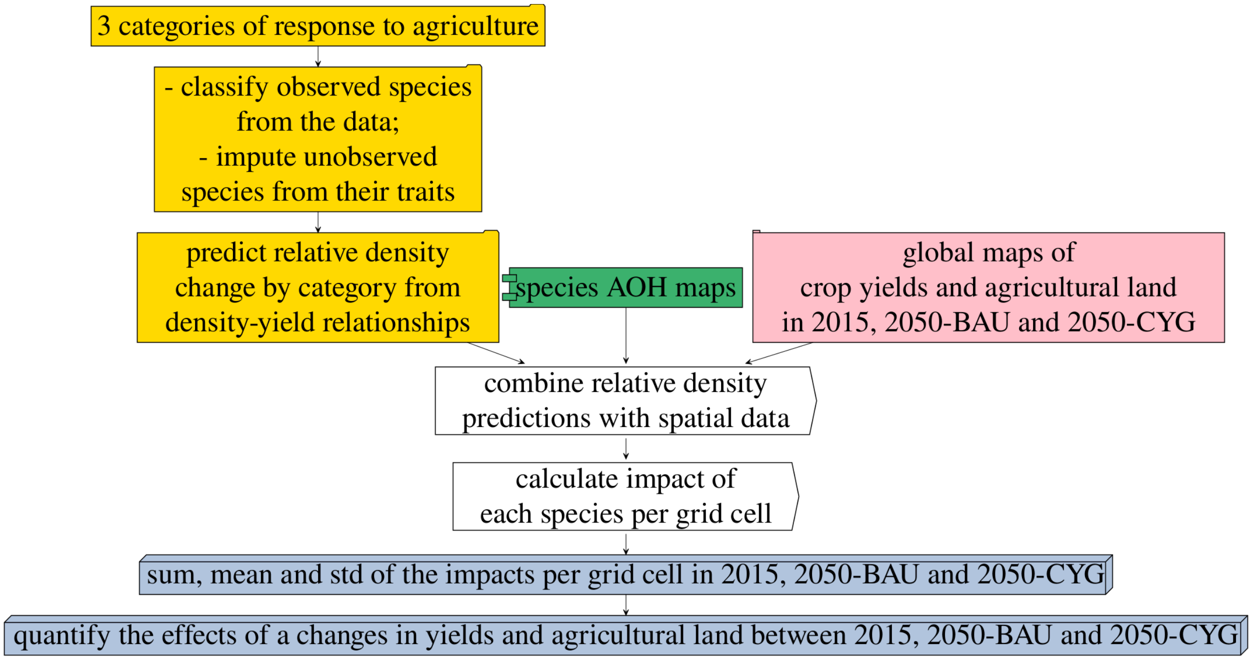

Figure S1: Flowchart showing the steps involved in the analysis and how we combined information from the agricultural projections, the vulnerability to agriculture (based on the interpolation from density-yield curves) to identify areas with higher impact from agriculture. The three scenarios considered—2015, 2050 business as usual (BAU), and 2050 closing yield gaps (CYG)—had different values of crop yields and agricultural land. Hence, different values of impacts were calculated under each scenario, and we conducted a quantitative comparison between them.

### Figure S2: Study sites location

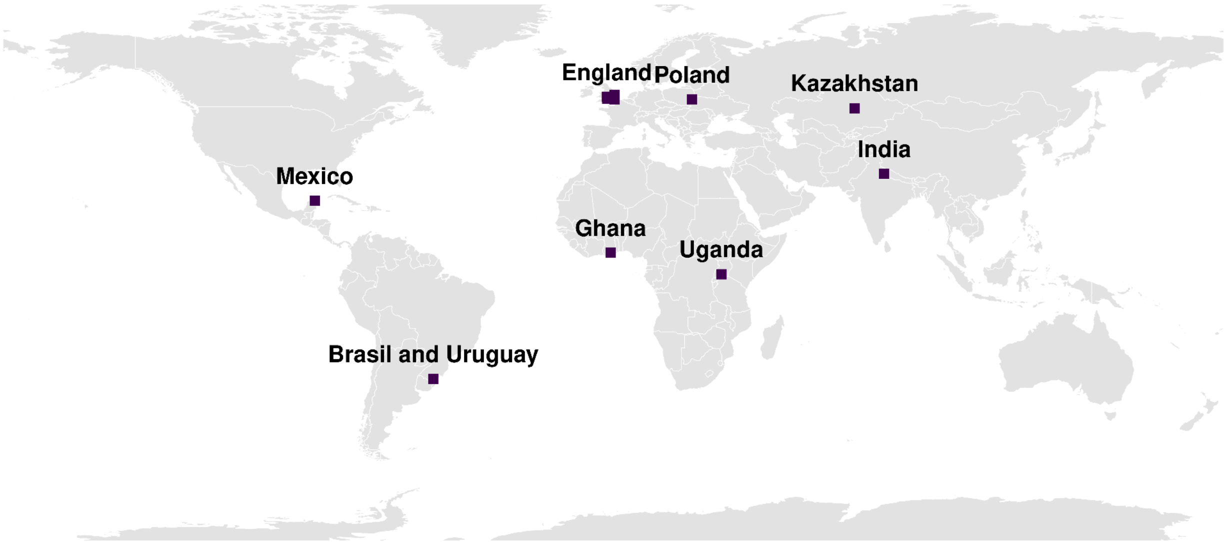

Figure S2: Location of the study sites: Mexico, Brasil and Uruguay, four sites in England, Poland, Ghana, Uganda, Kazakhstan and India.

#### Figure S3: Estimated mean crop yields by country

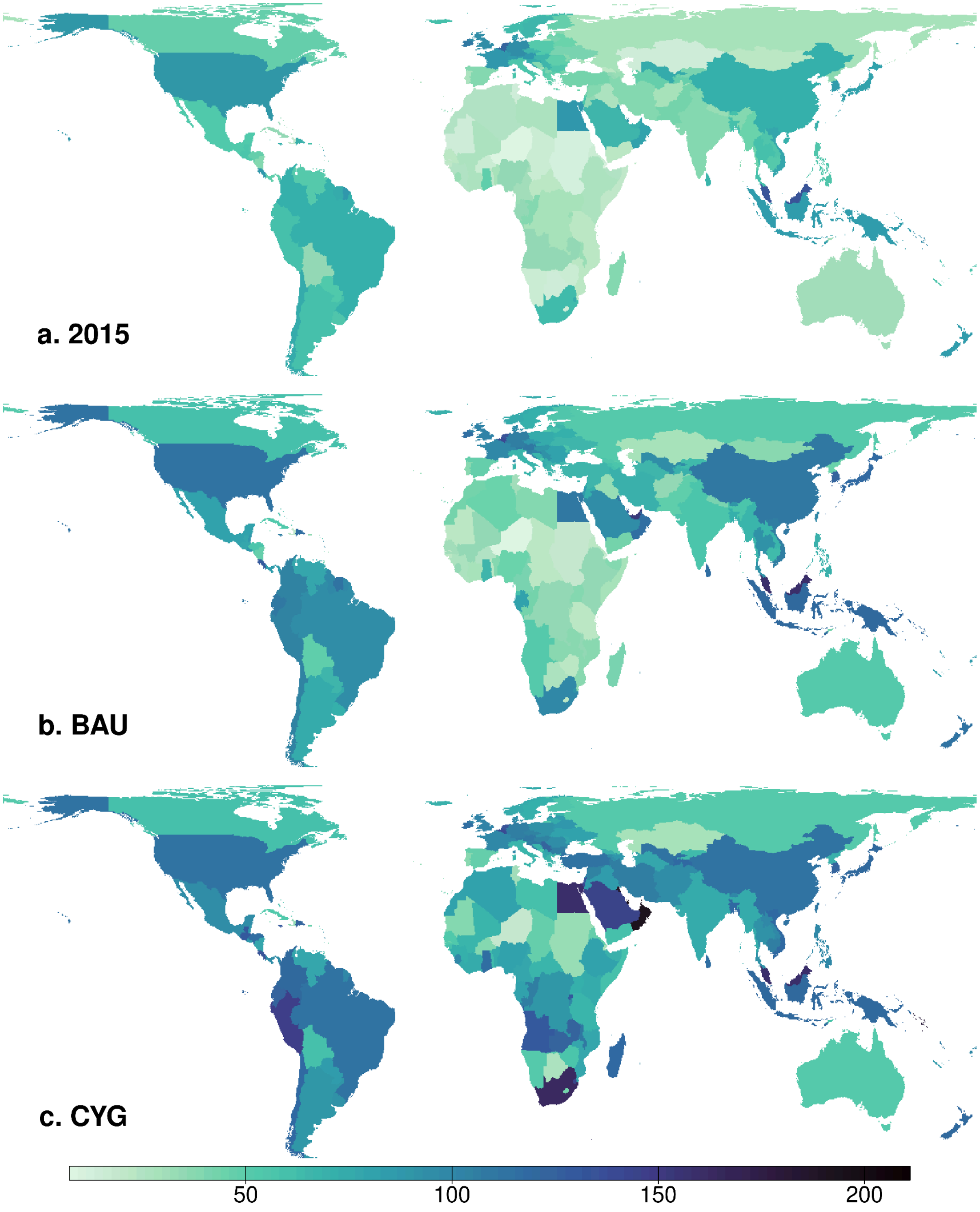

Figure S3: Map of mean crop yields per country in Gj/ha derived from Williams et al. (2021) set to the year 2015 (panel a. 2015), to the year 2050 under the business-as-usual scenario (panel b. BAU), and to the year 2050 with closing yield gaps (panel c. CYG)). Amongst the countries with lowest yields in 2015 were Niger (7.2 Gj/ha), Djibouti (10 Gj/ha) and the former Sudan (11.4 Gj/ha). Countries with the highest yields in 2015 included the Netherlands (122.2 Gj/ha), Belgium (125.7 Gj/h) and Malaysia (134.8 Gj/ha). Note that map polygons delineate study areas with homogeneous expected mean yields and do not necessarily depict national boundaries.

#### Figure S4: Estimated agricultural land

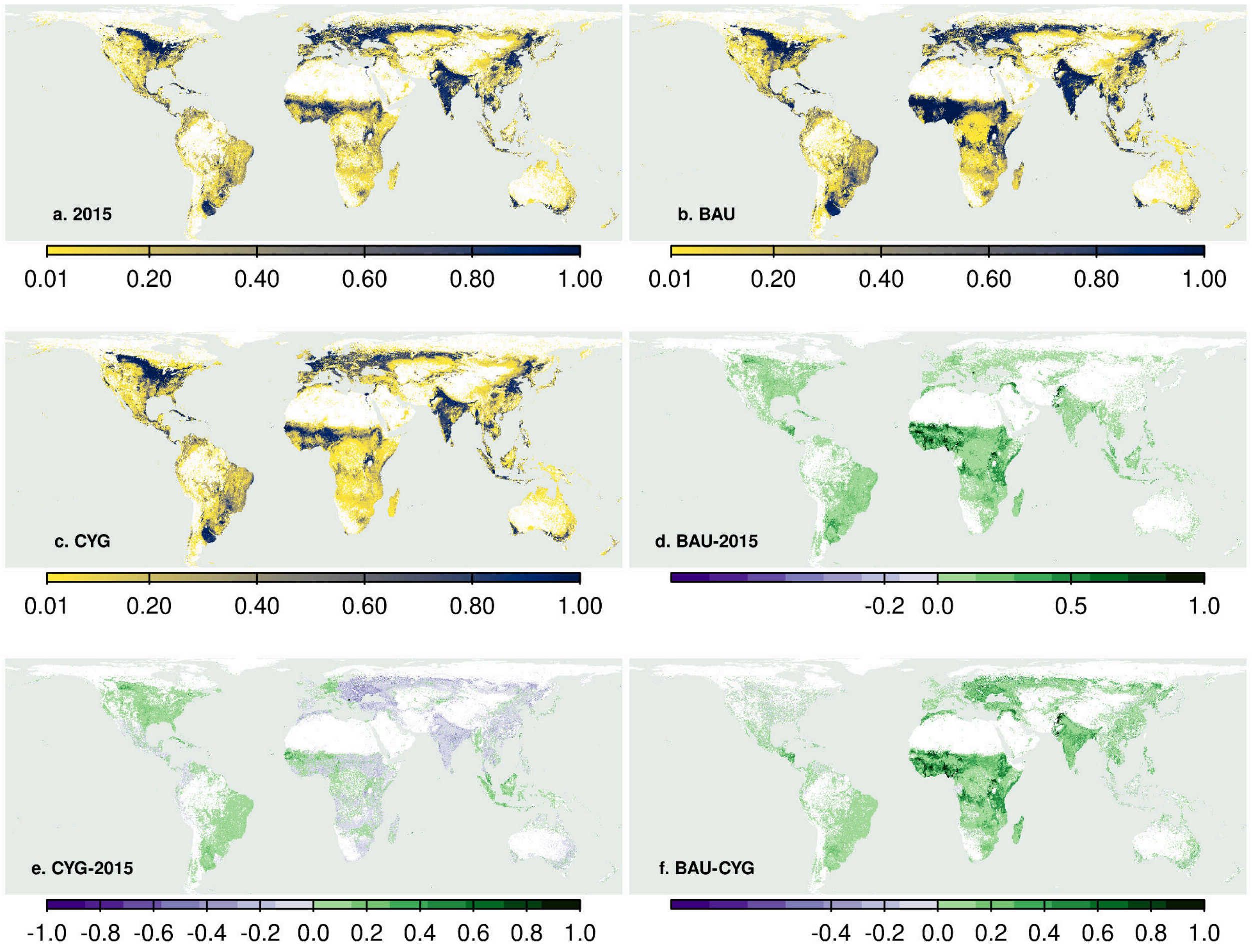

Figure S4: Map of the proportion of each pixel dedicated to agriculture derived from Williams et al. (2021) for the year 2015 (panel a. 2015), for the year 2050 under the business-as-usual scenario (panel b. BAU), and for the year 2050 within the close yield gap scenario (panel c. CYG)). We also plot the differences between the maps. Panel d. BAU-2015 represents the difference between BAU and 2015, panel e. CYG-2015 is the difference between CYG and 2015, and panel f. BAU-CYG is the difference between BAU and CYG. Green pixels indicate a positive difference (i.e., pixels where the first map has higher values than the second map). On the other hand, green pixels indicate a negative difference (i.e., the second map has higher values than the first). Notably, the difference between BAU minus 2015 (d. BAU-2015); and BAU minus CYG (f. BAU-CYG) are primarily green, indicating that larger percentages of pixels are dedicated to agriculture under BAU than 2015 or CYG. The difference between CYG minus 2015 is a mix of green and purple pixels, indicating that the agricultural land will expand in the green pixels and contract in the purple pixels under CYG when compared to 2015 (e. CYG-2015).

#### Figure S5: Curves relating species relative standardized density per hectare (ha) to agricultural yield for different response categories

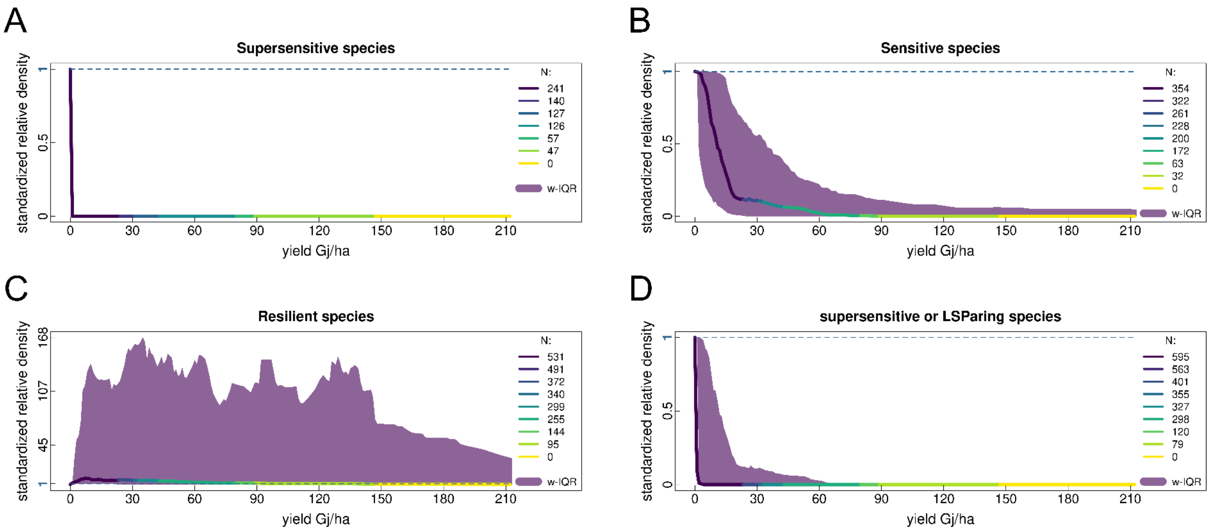

Figure S5: Curves relating species relative standardized density per hectare (ha) to agricultural yield (the production target in Gj/ha) of supersensitive (A), sensitive (B), and resilient (C) species. Panel (D) shows the curve for the simplified category (two-category models) combining supersensitive and Sensitive species. The line is the weighted median of the species in each category (species belonging to multiple sites have lower weights than species belonging to a single site). The colour of the line changes according to the number of species (N) with observational data that inform the line. The shaded purple areas represent the weighted interquartile ranges (w-IQR) around the median lines. The comparison between the median lines in panels A, B and C is shown in Figure 1 in the main text.

#### Figure S6: Comparison between the density-yield response categories and the tolerance to human pressure

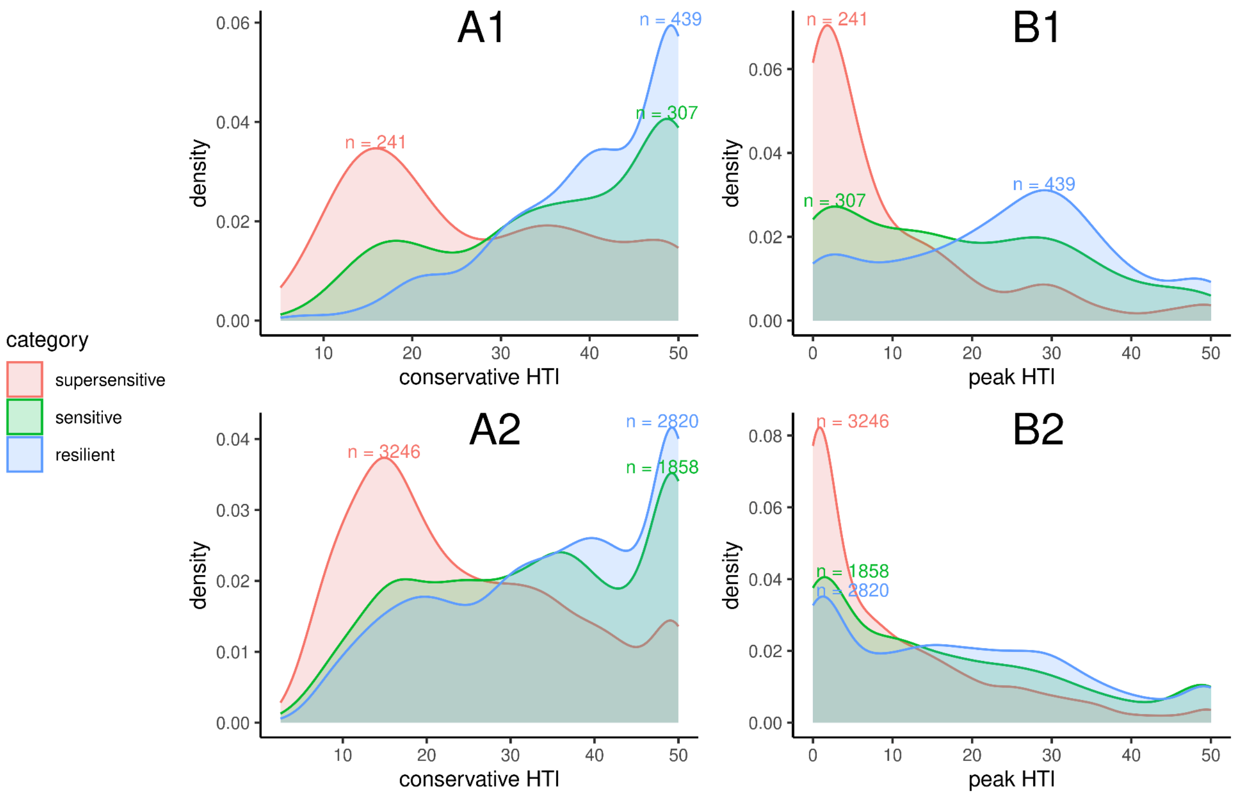

Figure S6: Comparison between our categories of density-yield response and the tolerance to human pressure (Human Tolerance Index or HTI; Marjakangas et al. 2024). Panels A1 and A2 show the comparison with the conservative HTI, while panels B1 and B2 with the peak HTI.

### Figure S7: Ternary plot

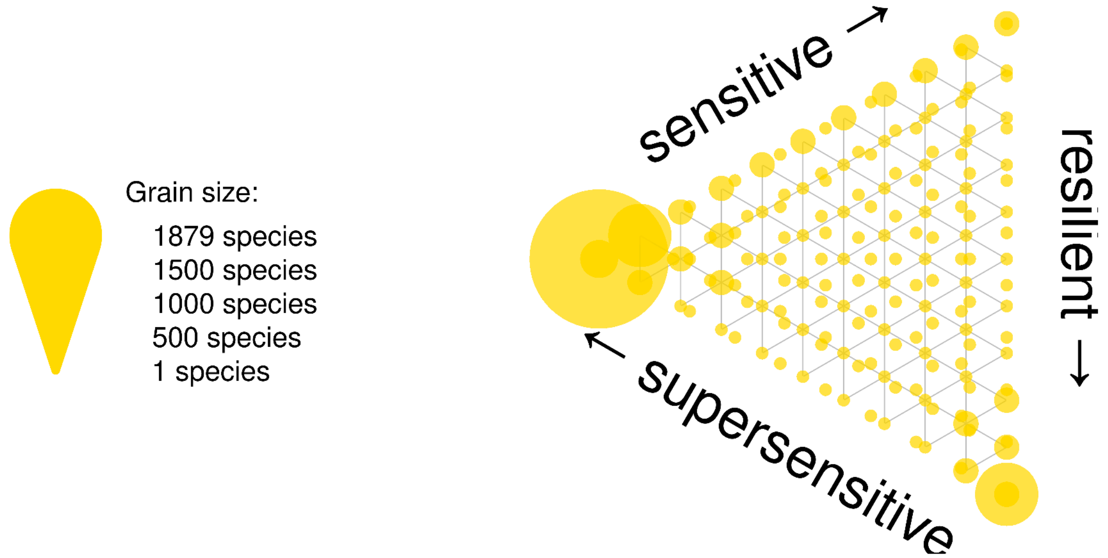

Figure S7: Ternary plot illustrating the uncertainty in the classification of all species (observed and predicted). The three vertices of the triangle represent species classified with a probability of 1 as supersensitive (left vertex), sensitive (top right vertex), and resilient (bottom vertex). Points within the triangle indicate specific compositions of probabilities of belonging to the three classes. The grain size of the yellow circles corresponds to the number of species located at the same position on the plot. For example, the largest circle represents 1,879 species classified with a probability of 1 as supersensitive, while the smallest circles typically represent fewer than 11 species with similar probabilities across the three classes. The circles at the sensitive and resilient vertices correspond to 280 and 766 species, respectively.

#### Figure S8: Mean impact in the three scenarios

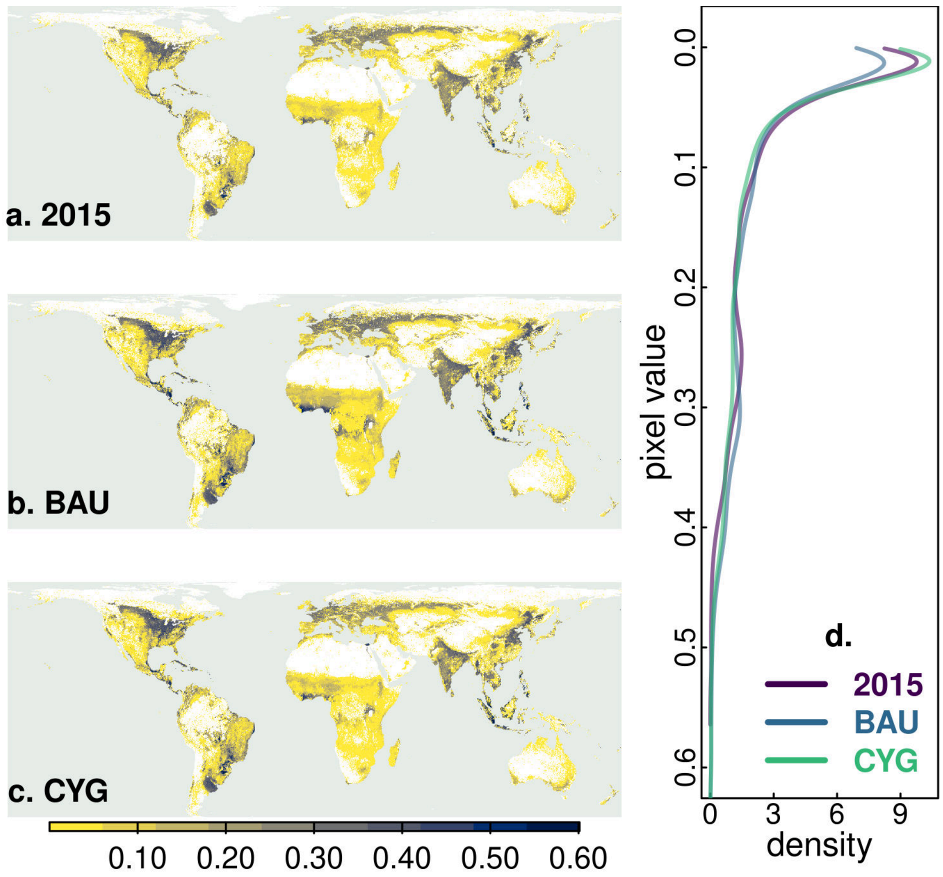

Figure S8: Maps of the mean impact of all species in a pixel in 2015 (panel a. 2015), under 2050 business as usual (panel b. BAU) and 2050 close yield gaps (panel c. CYG). In panel (d) we plot the density distribution of the pixel values corresponding to the three maps. Pixel values are capped to the 0.999 quantiles.

### Figure S9: Standard deviation of the mean impact

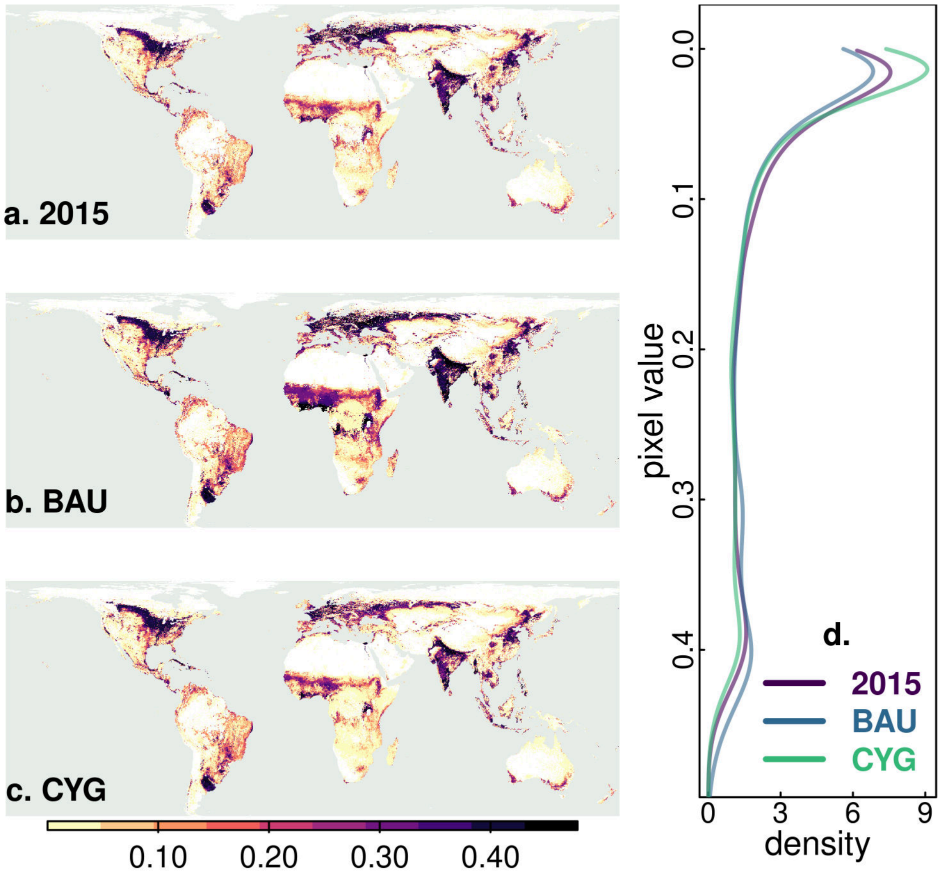

Figure S9: Map of the standard deviation around the mean impact (see Figure above) per pixel in the agriculture scenarios 2015 (panel a. 2015), 2050 business as usual (panel b. BAU) and 2050 close yield gap (panel c. CYG). In panel (d) we plot the density distribution of the pixel values corresponding to the three maps. Upper pixel values are truncated to the 0.999 quantile.

### Figure S10: Proportion of species with projected density declines

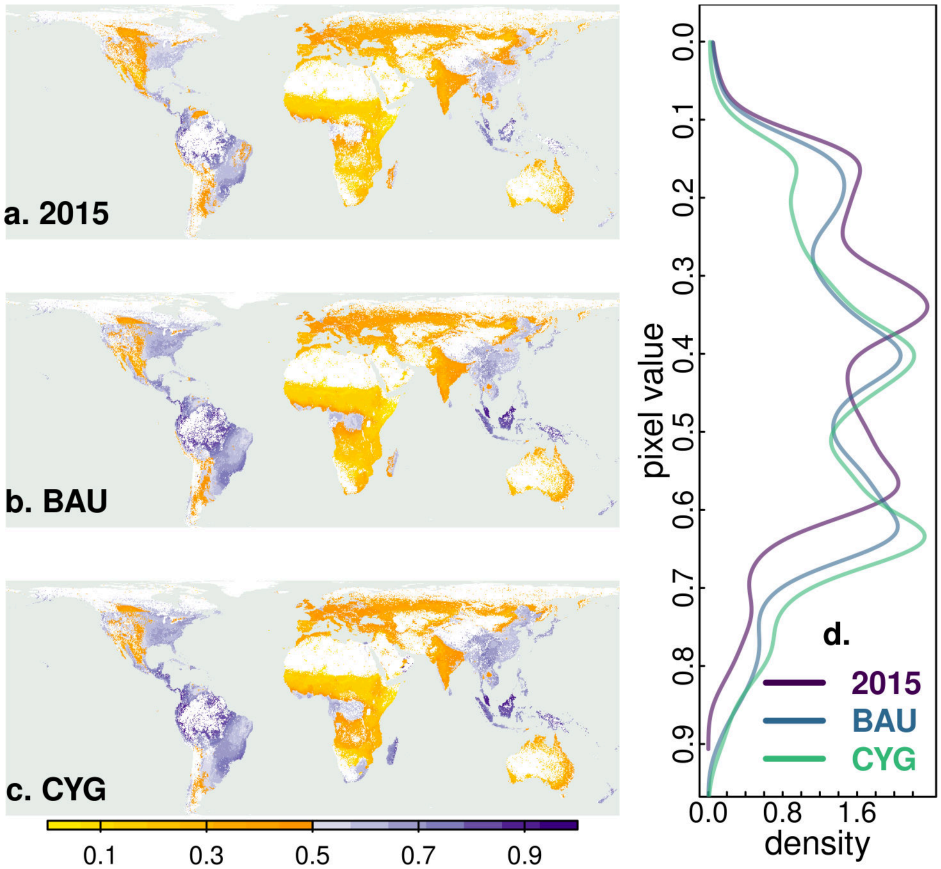

Figure S10: Map of the proportion of species with projected density declines per pixel in the agriculture scenarios 2015 (panel a. 2015), 2050 business as usual (panel b. BAU) and 2050 close yield gap (panel c. CYG). In panel (d), we plot the density distribution of the pixel values corresponding to the three maps.

### Figure S11: Interquartile range of the impact

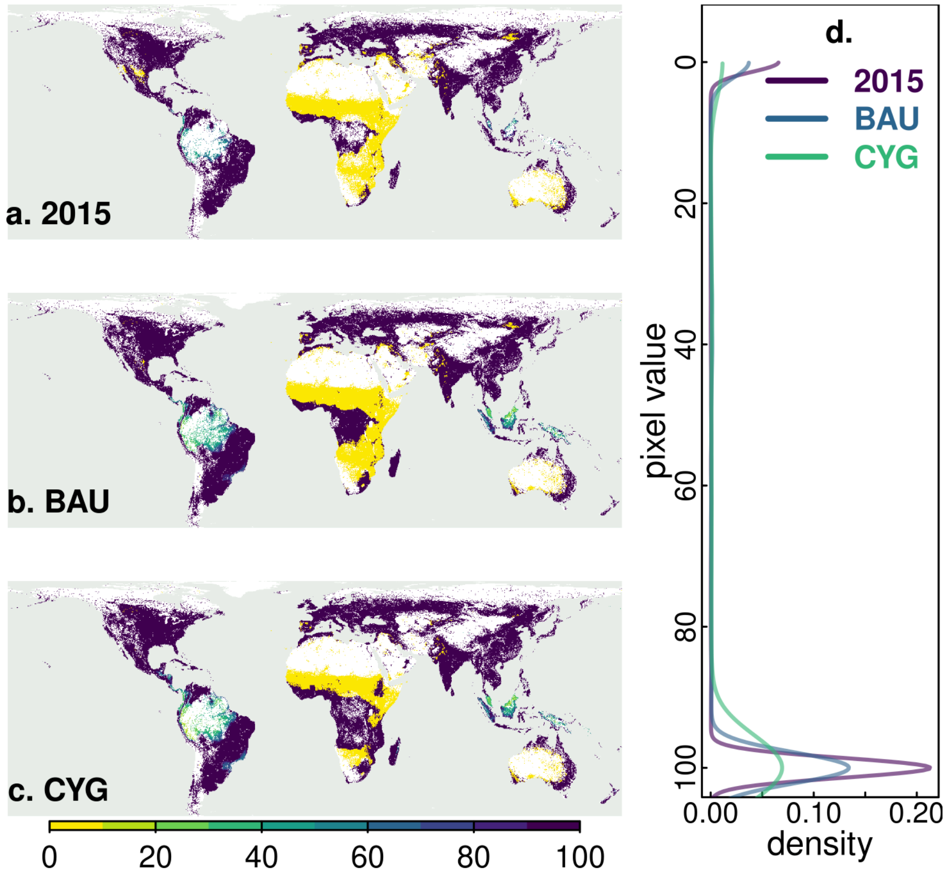

Figure S11: Map of the interquartile range, i.e., the distance between the 0.75 and 0.25 quartiles of the vulnerability of all species in a cell. Larger values indicate a larger distribution spread between species occupying the same cell.

#### Figure S12: Impact maps based on two density-yield response categories

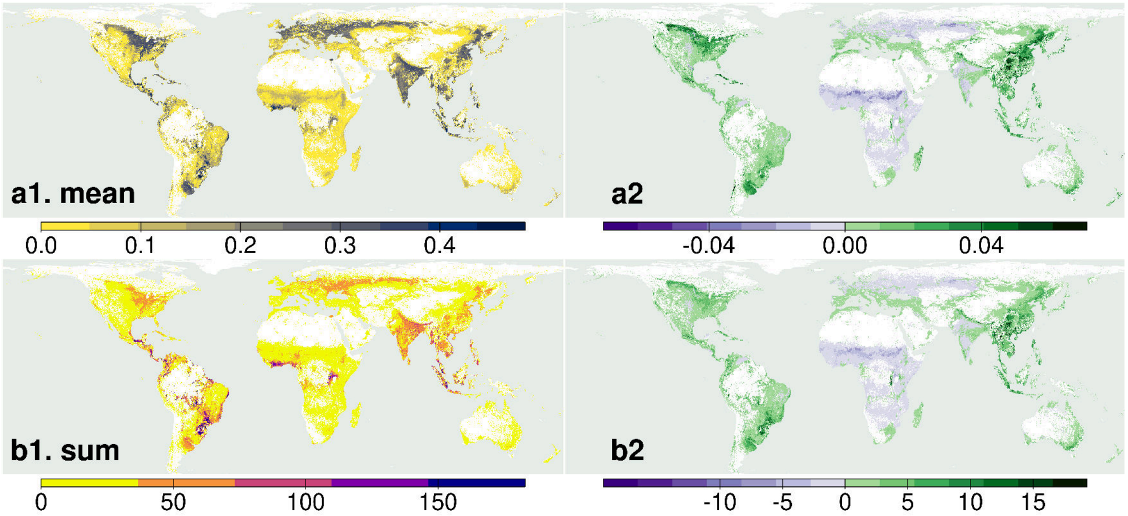

Figure S12: Maps of the mean impact by pixel (panel a1. mean), and sum of impacts of all species in a pixel (panel b1.sum), based on the 2015 national crop yields and crop land and derived from a two-category classification of species (disadvantaged, combining supersensitive and sensitive, and resilient species). The second column (panels a2 and b2) plots the difference between the values derived from the three-category maps and the two-category maps. Positive differences (green) suggest that the three-category maps overestimate the corresponding impact metric, negative (purple) suggest that the three-category maps underestimate impact. Values in the first column are capped at their 0.999 upper quantiles and those in the second column at both their lower (0.001) and upper (0.999) quantiles to reduce the influence of outliers.

### Figure S13: Impact maps based on globally threatened species only

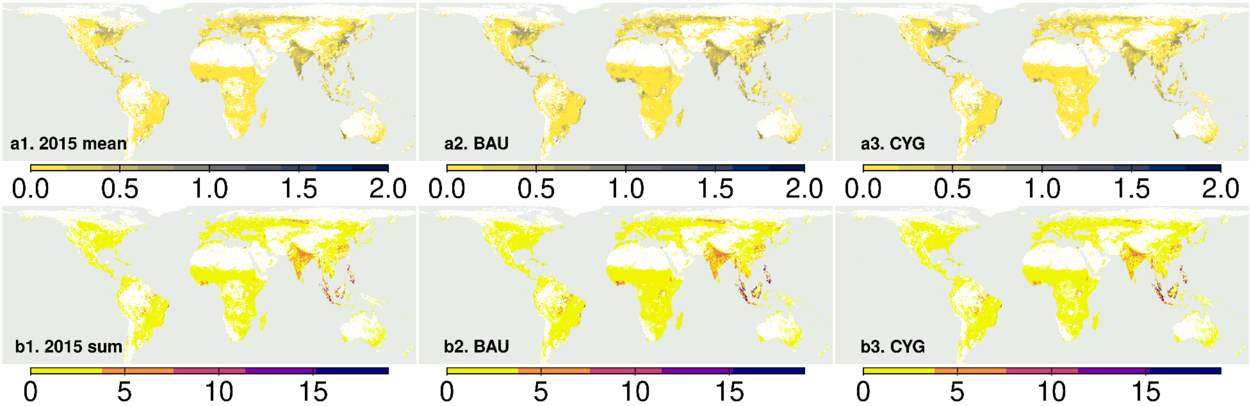

Figure S13: Maps of the mean impact (panels starting with A), and sum of impacts of all species in a pixel (panels starting with B), based on species listed as Vulnerable, Endangered or Critically Endangered on the IUCN Red List. We plot separately the three scenarios: 2015 (2015), 2050 business as usual (BAU) and 2050 close yield gaps (CYG). The small number of globally threatened species in some parts of the world causes extreme values in A. Values in all maps are capped at their 0.999 upper quantiles and 0.001 lower quantiles to reduce the influence of outliers.

#### Figure S14: Results without the correction with AOH size

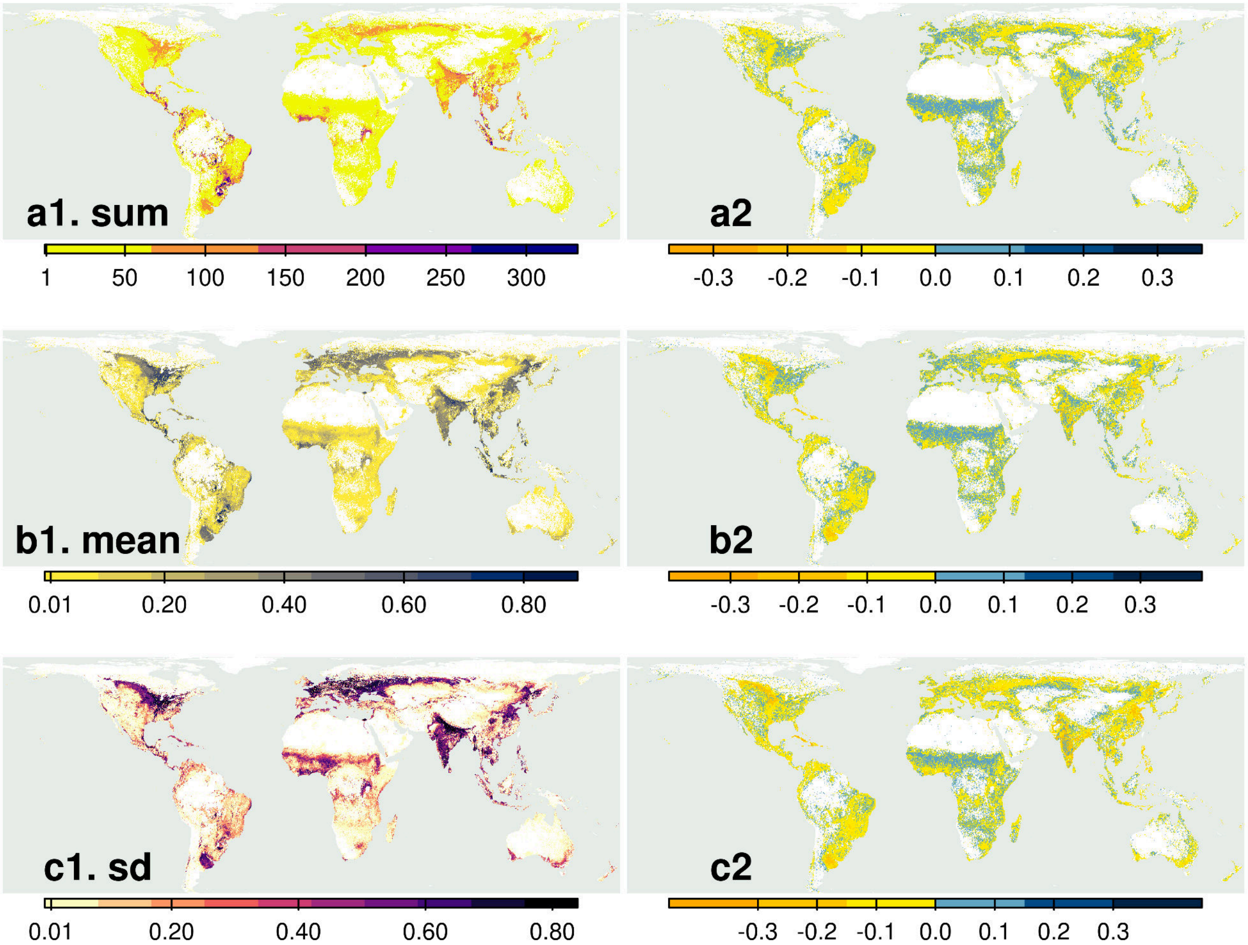

Figure S14: Maps illustrating the total impact of all species in a pixel (A), mean impact (B), and standard deviation (C) for the 2015 scenario, where the Area of Habitat (AOH) size adjustment was excluded in the calculation of species vulnerabilities. The right column shows the differences between these maps and the corresponding maps calculated with the AOH size adjustment, displayed in panels A-d (sum), B-d (mean), and C-d (standard deviation). In the right column, pixel values were standardised between 0 and 1 before calculating the differences. This standardisation was applied to preserve the relative importance of each pixel across scenarios, noting that the maps without the AOH size adjustment have larger values due to the lack of AOH size (a positive value between 1 and 100) in the denominator of the equation (Appendix S2). All values are capped at the 0.999 upper quantile to minimise the impact of outliers. The comparison suggests that while the absolute pixel values differ due to the distinct equations used in the two versions, the patterns in the maps with and without AOH size correction are similar.

### Figure S15: Histogram of cell values in 2015, BAU and CYG

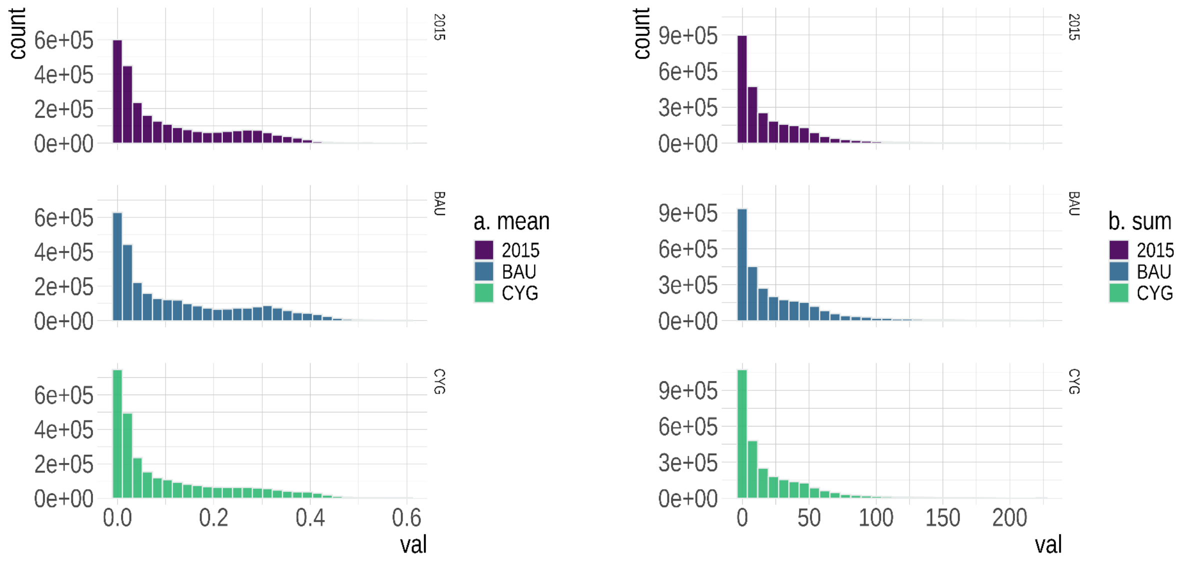

Figure S15: Histogram of the distribution of cell value globally for metric a (mean impact per cell) and for metric b (sum of impacts per cell) in the three scenarios: 2015 (panels a1. 2015 mean and b1. 2015 sum), BAU (a2. BAU and b2. BAU), and CYG (a3. CYG and b3. CYG)

#### Figure S16: Histogram of cell values in (BAU minus 2015) and (CYG minus 2015)

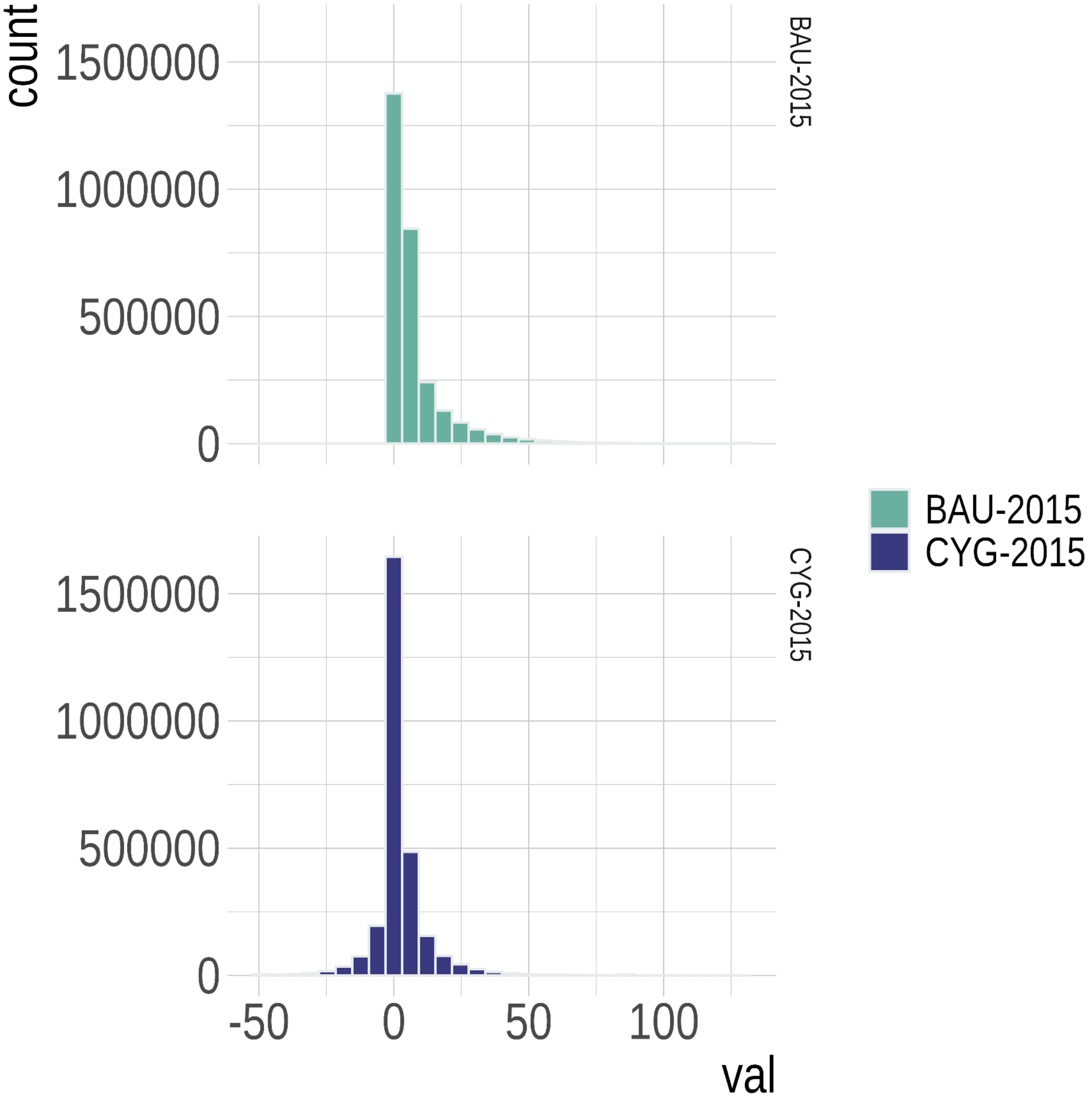

Figure S16: Histogram of the distribution of cell value globally for the sum of impacts per cell in the the difference between BAU minus 2015 (BAU-2015), and CYG minus 2015 (CYG-2015)

#### Figure S17: Histogram of cell values in (BAU minus CYG)

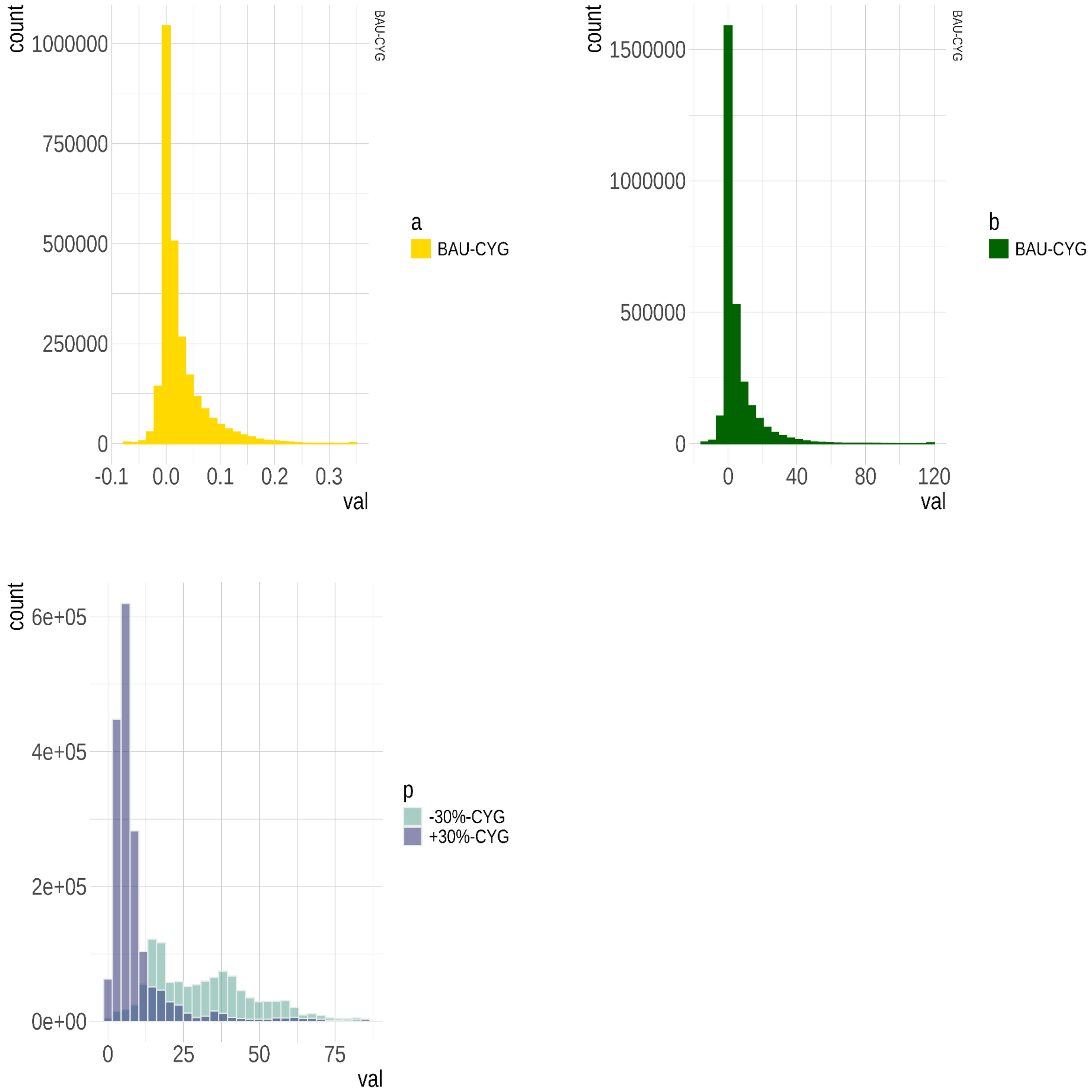

Figure S17: Histograms of the distribution of cell value globally of the difference between the BAU and CYG for metric A (mean impact per cell), B (sum of impacts per cell) and of the proportion of species showing a 30% increase (+30%-CYG) and a 30% decrease (-30%-CYG) in impact between the BAU and CYG (panel P).

### Table S1: Variables used for modelling density-yield response categories

Table S1: Predictors used in the multinomial classification model to predict the category of impact to agriculture for unobserved species. All variables are publicly available from different sources. We report the reference for each predictor (Tobias et al. 2022; Jetz et al. 2012; Bird et al. 2020;IUCN 2021-3), the type (continuous, dichotomous, or categorical), the units, and a short description. The description for the Avonet database was copy-pasted from Tobias et al. 2022. The values in the column predictor correspond to the column names used in the software implementation.

| predictor | reference | type | units | description |
|---|---|---|---|---|
| avonet__beak__depth | Tobias et al 2022 | continuous | mm | Depth of the beak at the anterior edge of the nostrils |
| avonet__beak__length__culmen | Tobias et al 2022 | continuous | mm | Length from the tip of the beak to the base of the skull |
| avonet__beak__length__nares | Tobias et al 2022 | continuous | mm | Length from the anterior edge of the nostrils to the tip of the beak |
| avonet__beak__width | Tobias et al 2022 | continuous | mm | Width of the beak at the anterior edge of the nostrils |
| avonet__habitat | Tobias et al 2022 | categorical |  | Desert (= drylands and other open arid habitats often sandy with very sparse vegetation)<br>Rock (= rocky substrate typically with no or very little vegetation including rocky outcrops rocky coastlines arid stony steppes rocky mountaintops and mountain slopes)<br>Grassland (= open dry to moist grass-dominated landscapes at all elevations)<br>Shrubland (= low stature bushy habitats included thornscrub thorny or arid savanna caatinga xerophytic shrubland and coastal scrub)<br>Woodland (= medium stature tree-dominated habitats including Acacia woodland riparian woodlands mangrove forests forest edges also more open parkland with scattered taller trees)<br>Forest (= tall tree-dominated vegetation with more or less closed canopy including palm forest)<br>Human modified (urban landscapes intensive agriculture gardens)<br>Wetland (= wide range of freshwater aquatic habitats including lakes marshes swamps and reedbeds)<br>Riverine (= associated with rivers and streams at all elevations)<br>Coastal (= intertidal zones within immediate vicinity of beaches estuaries brackish to salty marshes including mudflats lagoons alkaline wetlands coastal dunes and harbours)<br>Marine (= pelagic on sea near coasts including species in the intertidal zone on beaches and those pelagic species nesting near the sea on cliffs islets and islands). |

Table S1: Predictors used in the multinomial classification model to predict the category of impact to agriculture for unobserved species. All variables are publicly available from different sources. We report the reference for each predictor (Tobias et al. 2022; Jetz et al. 2012; Bird et al. 2020;IUCN 2021-3), the type (continuous, dichotomous, or categorical), the units, and a short description. The description for the Avonet database was copy-pasted from Tobias et al. 2022. The values in the column predictor correspond to the column names used in the software implementation.

| predictor | reference | type | units | description |
|---|---|---|---|---|
| avonet__habitat__density | Tobias et al 2022 | discrete | | 1 = Dense habitats. Species primarily lives in the lower or middle storey of forest or in dense thickets dense shrubland etc.<br>2 = Semi-open habitats. Species primarily lives in open shrubland scattered bushes parkland low dry or deciduous forest thorn forest.<br>3 = Open habitats. Species primarily lives in desert grassland open water low shrubs rocky habitats seashores cities. Also applies to species living mainly on top of forest canopy (i.e. mostly in the open) |
| avonet__hand__wing__index | Tobias et al 2022 | continuous | | 100*DK/Lw where DK is Kipp's distance and Lw is wing length (i.e. Kipp's distance corrected for wing size). Species average HWI differ from estimates in Sheard et al. (2020) because of much higher sampling of individuals in some species as well as taxonomic effects in the BirdLife list |
| avonet__kipps__distance | Tobias et al 2022 | continuous | mm | Length from the tip of the first secondary feather to the tip of the longest primary |
| avonet__primary__lifestyle | Tobias et al 2022 | categorical | | Aerial = species spends much of the time in flight and hunts or forages predominantly on the wing<br>Terrestrial = species spends majority of its time on the ground where it obtains food while either walking or hopping (note this includes species that also wade in water with their body raised above the water)<br>Insessorial = species spends much of the time perching above the ground either in branches of trees and other vegetation (i.e. arboreal) or on other raised substrates including rocks buildings posts and wires<br>Aquatic = species spends much of the time sitting on water and obtains food while afloat or when diving under the water's surface<br>Generalist = species has no primary lifestyle because it spends time in different lifestyle classes |
| avonet__secondary1 | Tobias et al 2022 | continuous | mm | Length from the carpal joint (bend of the wing) to the tip of the first secondary i.e. the outermost secondary adjacent to the innermost primary feather. Secondary1 is roughly equivalent to Wing length minus Kipp's distance (measured in a fully folded and flat wing) |
| avonet__tail__length | Tobias et al 2022 | continuous | mm | Distance between the tip of the longest rectrix and the point at which the two central rectrices protrude from the skin typically measured using a ruler inserted between the two central rectrices |
| avonet__tarsus__length | Tobias et al 2022 | continuous | mm | Length of the tarsus from the posterior notch between tibia and tarsus to the end of the last scale of acrotarsium (at the bend of the foot) |
| avonet__trophic__level | Tobias et al 2022 | categorical | | Herbivore = species obtaining at least 70% of food resources from plants/ Carnivore = species obtaining at least 70% of food resources by consuming live invertebrate or vertebrate animals / Scavenger = species obtaining at least 70% of food resources from carrion or refuse/ Omnivore = species obtaining resources from multiple trophic level in roughly equal proportion |

Table S1: Predictors used in the multinomial classification model to predict the category of impact to agriculture for unobserved species. All variables are publicly available from different sources. We report the reference for each predictor (Tobias et al. 2022; Jetz et al. 2012; Bird et al. 2020;IUCN 2021-3), the type (continuous, dichotomous, or categorical), the units, and a short description. The description for the Avonet database was copy-pasted from Tobias et al. 2022. The values in the column predictor correspond to the column names used in the software implementation.

| predictor | reference | type | units | description |
|---|---|---|---|---|
| avonet__trophic__niche | Tobias et al 2022 | categorical | | Frugivore = species obtaining at least 60% of food resources from fruit<br>Granivore = species obtaining at least 60% of food resources from seeds or nuts<br>Nectarivore = species obtaining at least 60% of food resources from nectar<br>Herbivore = species obtaining at least 60% of food resources from other plant materials in non-aquatic systems including leaves buds whole flowers etc.<br>Herbivore aquatic = species obtaining at least 60% of food resources from plant materials in aquatic systems including algae and aquatic plant leaves<br>Invertivore = species obtaining at least 60% of food resources from invertebrates in terrestrial systems including insects worms arachnids etc.<br>Vertivore = species obtaining at least 60% of food resources from vertebrate animals in terrestrial systems including mammals birds reptiles etc.<br>Aquatic Predator = species obtaining at least 60% of food resources from vertebrate and invertebrate animals in aquatic systems including fish crustacea molluscs etc<br>Scavenger = species obtaining at least 60% of food resources from carrion offal or refuse Omnivore = Species using multiple niches within or across trophic levels in relatively equal proportions |
| avonet__wil__mass__grams | Tobias et al 2022 | continuous | gram | Body mass given as species average (incorporating both male and female body mass) |
| avonet__wing__length | Tobias et al 2022 | continuous | mm | Length from the carpal joint (bend of the wing) to the tip of the longest primary on the unflattened wing |
| bird__age__at__first__breeding | Bird et al 2020 | continuous | years | the average age of parents of the current cohort |
| bird__longevity | Bird et al 2020 | continuous | years | maximum longevity |
| bird__survival | Bird et al 2020 | continuous | years | annual adult survival |
| num__pixels__suitable | IUCN 2021-3 | continuous | | number of suitable pixels with the area of habitat (AOH) map |
| phylogeny__distance__from__root | Jetz et al 2012 | continuous | | number of phylogenetic nodes from the last common ancestor |
| range__map__size | IUCN 2021-3 | continuous | | number of suitable and unsuitable pixels in the AOH map |
| rl__category | IUCN 2021-3 | continuous | | red list category |
| rl__count__hab__level__1 | IUCN 2021-3 | continuous | | total number of suitable habitats at level 1 (see IUCN Habitat Classification Scheme for the difference between level 1 and 2) |
| rl__count__hab__level__2 | IUCN 2021-3 | continuous | | total number of suitable habitats at level 2 (see IUCN Habitat Classification Scheme for the difference between level 1 and 2) |
| rl__elevation__range | IUCN 2021-3 | continuous | | difference between the maximum and minimum altitude range |

Table S1: Predictors used in the multinomial classification model to predict the category of impact to agriculture for unobserved species. All variables are publicly available from different sources. We report the reference for each predictor (Tobias et al. 2022; Jetz et al. 2012; Bird et al. 2020;IUCN 2021-3), the type (continuous, dichotomous, or categorical), the units, and a short description. The description for the Avonet database was copy-pasted from Tobias et al. 2022. The values in the column predictor correspond to the column names used in the software implementation.

| predictor | reference | type | units | description |
|---|---|---|---|---|
| rl__eoo__mean | IUCN 2021-3 | continuous | | species extent of occurrence (mean) |
| rl__freshwater | IUCN 2021-3 | binary | | 1 if the bird is riparian 0 otherwise |
| rl__generation__length__range | IUCN 2021-3 | continuous | | generation length |
| rl__habitat__code__1 [__2 up to __15__imp] | IUCN 2021-3 | dicothomous | | suitability value for each redlist habitat code (the number correspond to the IUCN habitat classification scheme level 1) |
| rl__habitat__code__1__imp [__2__imp up to __15__imp] | IUCN 2021-3 | continuous | | combination of habitat suitability and habitat importance: 0 = not suitable/ 1 = suitability is marginal or unknown/ 2 = suitable but not major/ 3 = suitable and major |
| rl__marine | IUCN 2021-3 | binary | | 1 if the bird is marine 0 otherwise |
| rl__mid__point__elevation | IUCN 2021-3 | continuous | | mid point between the min and max altitudinal range |
| rl__movement__patterns__pattern | IUCN 2021-3 | categorical | | migrant = migrant species \| not__migrant = resident species\| Nomadic = nomadic species\|altitidinal migrant = altitudinal migrant |
| rl__order__name | IUCN 2021-3 | categorical | | order name (ACCIPITRIFORMES ANSERIFORMES BUCEROTIFORMES CAPRIMULGIFORMES CHARADRIIFORMES CICONIIFORMES COLUMBIFORMES CORACIIFORMES CUCULIFORMES FALCONIFORMES GALLIFORMES GRUIFORMES MUSOPHAGIFORMES PASSERIFORMES PELECANIFORMES PICIFORMES PODICIPEDIFORMES PSITTACIFORMES rare STRIGIFORMES STRUTHIONIFORMES SULIFORMES TROGONIFORMES. The rare order combines orders with only a few species and not represented in the observed dataset: CARIAMIFORMES CATHARTIFORMES COLIIFORMES EURYPYGIFORMES LEPTOSOMIFORMES MESITORNITHIFORMES OPISTHOCOMIFORMES OTIDIFORMES PHOENICOPTERIFORMES PTEROCLIFORMES"" ) |
| rl__population__size__mean | IUCN 2021-3 | continuous | | mean population size |
| rl__population__size__range__derived | IUCN 2021-3 | continuous | | difference between the maximum and minimum population size |
| rl__severe__fragmentation__is__fragmented | IUCN 2021-3 | binary | | 1 the habitat is heavily fragmented 0 otherwise |

### Table S2: User-defined model settings for the classification model

Table S2: Custom-settings used to run the multinomial classification algorithm with 50-folds cross-validation and Lasso-regularization in h2o.ai (please read the h2o.ai documentation for further details)

| **parameter** | **value** |
|---|---|
| nfolds | 50 |
| stopping_rounds | 5 |
| stopping_tolerance | 0.001 |
| stopping_metric | mean_per_class_error |
| distribution | multinomial |
| use_all_factor_levels | TRUE |
| standardize | TRUE |
| missing_values_handling | MeanImputation |
| variable_importances | TRUE |
| epochs | 1000 |
| loss | CrossEntropy |
| activation | Maxout |
| l1 | 0.001 |
| l2 | 0 |
| adaptive_rate | TRUE |
| categorical_encoding | OneHotInternal |
| reproducible | TRUE |
| verbose | TRUE |
| keep_cross_validation_predictions | TRUE |
| keep_cross_validation_fold_assignment | TRUE |
| shuffle_training_data | TRUE |
| seed | 1234 |

### Table S3: Country-wise validation of species classification

Table S3: Model performance when predicting species response to agriculture splitting the observed dataset into train (~80 percent) and validation (~20 percent) sets by country. For example, when we predicted Ghana and Uganda (see column country to predict), we included all the other countries in the test set and used the data from Ghana and Uganda for validation. For each separate model, we report the mean per class error, the log loss, the mean squared error, and the percentage of the data used for training.

| mean per class error | logloss | mean squared error | root-mean-square error | country to predict | percentage of data used for training |
|---:|---:|---:|---:|---|---:|
| 0.46 | 1.01 | 0.32 | 0.56 | Uganda | 80 |
| 0.55 | 3.04 | 0.46 | 0.68 | England | 75 |
| 0.65 | 2.05 | 0.53 | 0.72 | Brasil Mexico | 83 |
| 0.54 | 1.98 | 0.43 | 0.65 | Kazakhstan India Poland | 75 |
| 0.46 | 1.43 | 0.37 | 0.61 | Ghana Uganda | 68 |

### Table S4: The importance of the different predictors in classifying species to density-yield response categories

Table S4: The importance of the different predictors in classifying a species in a category of impact to agriculture.

| variable | relative__importa | scaled__importar | percentage |
|---|---|---|---|
| rl__order__name.BUCEROTIFORMES | 1.0000 | 1.0000 | 0.0103 |
| rl__order__name.PASSERIFORMES | 0.9999 | 0.9999 | 0.0103 |
| rl__habitat__code__5 | 0.9741 | 0.9741 | 0.0100 |
| rl__order__name.GALLIFORMES | 0.9326 | 0.9326 | 0.0096 |
| rl__order__name.CHARADRIIFORMES | 0.9318 | 0.9318 | 0.0096 |
| rl__population__size__range__derived | 0.9263 | 0.9263 | 0.0095 |
| rl__order__name.PODICIPEDIFORMES | 0.9236 | 0.9236 | 0.0095 |
| rl__order__name.PELECANIFORMES | 0.9189 | 0.9189 | 0.0095 |
| rl__order__name.GRUIFORMES | 0.9159 | 0.9159 | 0.0094 |
| rl__order__name.FALCONIFORMES | 0.9059 | 0.9059 | 0.0093 |
| rl__order__name.MUSOPHAGIFORMES | 0.9048 | 0.9048 | 0.0093 |
| rl__order__name.COLUMBIFORMES | 0.9017 | 0.9017 | 0.0093 |
| rl__habitat__code__15__imp | 0.8997 | 0.8997 | 0.0093 |
| rl__order__name.CUCULIFORMES | 0.8941 | 0.8941 | 0.0092 |
| rl__order__name.ACCIPITRIFORMES | 0.8906 | 0.8906 | 0.0092 |
| rl__order__name.PICIFORMES | 0.8849 | 0.8849 | 0.0091 |
| rl__order__name.CAPRIMULGIFORMES | 0.8827 | 0.8827 | 0.0091 |
| avonet__trophic__niche.Vertivore | 0.8795 | 0.8795 | 0.0091 |
| avonet__primary__lifestyle.Insessorial | 0.8768 | 0.8768 | 0.0090 |
| rl__habitat__code__14__imp | 0.8766 | 0.8766 | 0.0090 |
| rl__order__name.CICONIIFORMES | 0.8706 | 0.8706 | 0.0090 |
| rl__order__name.CORACIIFORMES | 0.8663 | 0.8663 | 0.0089 |
| avonet__trophic__niche.Omnivore | 0.8659 | 0.8659 | 0.0089 |
| rl__order__name.ANSERIFORMES | 0.8653 | 0.8653 | 0.0089 |
| rl__habitat__code__12__imp | 0.8637 | 0.8637 | 0.0089 |
| rl__order__name.SULIFORMES | 0.8549 | 0.8549 | 0.0088 |
| rl__habitat__code__6__imp | 0.8544 | 0.8544 | 0.0088 |
| avonet__trophic__niche.Nectarivore | 0.8476 | 0.8476 | 0.0087 |
| tot__aoh__size | 0.8456 | 0.8456 | 0.0087 |
| avonet__habitat.Riverine | 0.8445 | 0.8445 | 0.0087 |

Table S4: The importance of the different predictors in classifying a species in a category of impact to agriculture.

| variable | relative_importa | scaled_importar | percentage |
|---|---|---|---|
| avonet_habitat_density | 0.8422 | 0.8422 | 0.0087 |
| avonet_trophic_niche.Scavenger | 0.8418 | 0.8418 | 0.0087 |
| avonet_primary_lifestyle.Generalist | 0.8417 | 0.8417 | 0.0087 |
| avonet_trophic_niche.Herbivore aquatic | 0.8399 | 0.8399 | 0.0086 |
| rl_habitat_code_9_imp | 0.8389 | 0.8389 | 0.0086 |
| rl_habitat_code_5_imp | 0.8386 | 0.8386 | 0.0086 |
| avonet_trophic_niche.Herbivore terrestrial | 0.8363 | 0.8363 | 0.0086 |
| rl_category.VU | 0.8344 | 0.8344 | 0.0086 |
| avonet_trophic_niche.Frugivore | 0.8342 | 0.8342 | 0.0086 |
| avonet_habitat.Grassland | 0.8325 | 0.8325 | 0.0086 |
| avonet_trophic_niche.Invertivore | 0.8325 | 0.8325 | 0.0086 |
| avonet_habitat.Woodland | 0.8321 | 0.8321 | 0.0086 |
| rl_habitat_code_1_imp | 0.8307 | 0.8307 | 0.0085 |
| rl_habitat_code_14 | 0.8303 | 0.8303 | 0.0085 |
| avonet_habitat.Desert | 0.8302 | 0.8302 | 0.0085 |
| avonet_tail_length | 0.8275 | 0.8275 | 0.0085 |
| num_pixels_suitable | 0.8259 | 0.8259 | 0.0085 |
| avonet_habitat.Human Modified | 0.8187 | 0.8187 | 0.0084 |
| avonet_trophic_niche.Aquatic predator | 0.8175 | 0.8175 | 0.0084 |
| rl_habitat_code_13_imp | 0.8170 | 0.8170 | 0.0084 |
| rl_movement_patterns_pattern.migrant | 0.8154 | 0.8154 | 0.0084 |
| rl_habitat_code_2_imp | 0.8148 | 0.8148 | 0.0084 |
| rl_habitat_code_2 | 0.8134 | 0.8134 | 0.0084 |
| rl_habitat_code_13 | 0.8119 | 0.8119 | 0.0084 |
| rl_habitat_code_15 | 0.8118 | 0.8118 | 0.0084 |
| avonet_habitat.Rock | 0.8103 | 0.8103 | 0.0083 |
| avonet_kipps_distance | 0.8094 | 0.8094 | 0.0083 |
| avonet_trophic_level.Omnivore | 0.8090 | 0.8090 | 0.0083 |
| rl_habitat_code_12 | 0.8081 | 0.8081 | 0.0083 |
| rl_movement_patterns_pattern.Nomadic | 0.8077 | 0.8077 | 0.0083 |
| avonet_wil_mass_grams | 0.8075 | 0.8075 | 0.0083 |
| rl_generation_length_range | 0.8046 | 0.8046 | 0.0083 |
| avonet_trophic_level.Herbivore | 0.8043 | 0.8043 | 0.0083 |

Table S4: The importance of the different predictors in classifying a species in a category of impact to agriculture.

| variable | relative_importa | scaled_importar | percentage |
|---|---|---|---|
| rl_order_name.PSITTACIFORMES | 0.8031 | 0.8031 | 0.0083 |
| avonet_beak_depth | 0.8031 | 0.8031 | 0.0083 |
| rl_habitat_code_6 | 0.8006 | 0.8006 | 0.0082 |
| avonet_habitat.Coastal | 0.8003 | 0.8003 | 0.0082 |
| phylogeny_distance_from_root | 0.7990 | 0.7990 | 0.0082 |
| avonet_primary_lifestyle.Terrestrial | 0.7975 | 0.7975 | 0.0082 |
| seasonality_aoh.missing(NA) | 0.7952 | 0.7952 | 0.0082 |
| bird_age_at_first_breeding | 0.7946 | 0.7946 | 0.0082 |
| rl_habitat_code_4 | 0.7923 | 0.7923 | 0.0082 |
| rl_habitat_code_1 | 0.7920 | 0.7920 | 0.0082 |
| rl_category.EN | 0.7905 | 0.7905 | 0.0081 |
| rl_order_name.rare | 0.7886 | 0.7886 | 0.0081 |
| avonet_primary_lifestyle.Aerial | 0.7882 | 0.7882 | 0.0081 |
| rl_habitat_code_8_imp | 0.7878 | 0.7878 | 0.0081 |
| rl_category.CR | 0.7877 | 0.7877 | 0.0081 |
| avonet_habitat.Wetland | 0.7850 | 0.7850 | 0.0081 |
| rl_movement_patterns_pattern.not_migrant | 0.7841 | 0.7841 | 0.0081 |
| rl_marine | 0.7833 | 0.7833 | 0.0081 |
| avonet_secondary1 | 0.7823 | 0.7823 | 0.0081 |
| rl_order_name.STRUTHIONIFORMES | 0.7819 | 0.7819 | 0.0080 |
| rl_order_name.TROGONIFORMES | 0.7789 | 0.7789 | 0.0080 |
| avonet_wing_length | 0.7775 | 0.7775 | 0.0080 |
| bird_longevity | 0.7773 | 0.7773 | 0.0080 |
| rl_habitat_code_7_imp | 0.7763 | 0.7763 | 0.0080 |
| avonet_hand_wing_index | 0.7757 | 0.7757 | 0.0080 |
| rl_eoo_mean | 0.7749 | 0.7749 | 0.0080 |
| avonet_habitat.Shrubland | 0.7729 | 0.7729 | 0.0080 |
| rl_freshwater | 0.7713 | 0.7713 | 0.0079 |
| rl_count_hab_level_1 | 0.7675 | 0.7675 | 0.0079 |
| avonet_beak_length_culmen | 0.7669 | 0.7669 | 0.0079 |
| rl_habitat_code_4_imp | 0.7635 | 0.7635 | 0.0079 |
| rl_order_name.STRIGIFORMES | 0.7628 | 0.7628 | 0.0079 |
| avonet_tarsus_length | 0.7529 | 0.7529 | 0.0077 |

Table S4: The importance of the different predictors in classifying a species in a category of impact to agriculture.

| variable | relative_importa | scaled_importar | percentage |
|---|---|---|---|
| rl_habitat_code_3_imp | 0.7475 | 0.7475 | 0.0077 |
| rl_movement_patterns_pattern.Altitudinal Migrant | 0.7455 | 0.7455 | 0.0077 |
| rl_category.LC | 0.7417 | 0.7417 | 0.0076 |
| seasonality_aoh.Resident | 0.7413 | 0.7413 | 0.0076 |
| rl_habitat_code_9 | 0.7411 | 0.7411 | 0.0076 |
| rl_population_size_mean | 0.7372 | 0.7372 | 0.0076 |
| range_map_size | 0.7353 | 0.7353 | 0.0076 |
| rl_category.NT | 0.7329 | 0.7329 | 0.0075 |
| bird_survival | 0.7310 | 0.7310 | 0.0075 |
| rl_habitat_code_8 | 0.7307 | 0.7307 | 0.0075 |
| rl_mid_point_elevation | 0.7273 | 0.7273 | 0.0075 |
| avonet_habitat.Marine | 0.7237 | 0.7237 | 0.0074 |
| rl_count_hab_level_2 | 0.7230 | 0.7230 | 0.0074 |
| rl_habitat_code_3 | 0.7223 | 0.7223 | 0.0074 |
| avonet_primary_lifestyle.Aquatic | 0.7218 | 0.7218 | 0.0074 |
| avonet_trophic_level.Scavenger | 0.7180 | 0.7180 | 0.0074 |
| rl_elevation_range | 0.7178 | 0.7178 | 0.0074 |
| avonet_trophic_niche.Granivore | 0.7146 | 0.7146 | 0.0074 |
| avonet_trophic_level.Carnivore | 0.7121 | 0.7121 | 0.0073 |
| seasonality_aoh.migrant | 0.7101 | 0.7101 | 0.0073 |
| avonet_habitat.Forest | 0.6992 | 0.6992 | 0.0072 |
| rl_habitat_code_7 | 0.6751 | 0.6751 | 0.0069 |
| avonet_beak_width | 0.6667 | 0.6667 | 0.0069 |
| avonet_beak_length_nares | 0.6350 | 0.6350 | 0.0065 |

### Table S5: Classification into two-categories of response to agriculture

Table S5: Model performance and confusion matrix of the species classification into two categories (disadvantaged and resilient) of impact to agriculture.

| | **mean per class error** | **logloss** | **mean squared error** | **root-mean-square error** |
|---|---|---|---|---|
| | 0.31 | 0.97 | 0.23 | 0.48 |
| **Confusion matrix** | | **Imputed** | | |
| **Observed** | **disadvantaged** | **resilient** | **Error rate** | **Success rate** |
| **Disadvantaged** | **388** | 331 | 331/719 | 0.54 |
| **Resilient** | 88 | **495** | 88/583 | 0.85 |
| **Totals** | 476 | 826 | 419/1302 | **0.68** |

### Table S6: Country-wise results of species classification into three categories of response to agriculture

Table S6: Confusion matrices associated with the multinomial classification model (three categories) splitting the dataset into train and validation sets by country combinations. In the last row, we report the average error rate of 0.49. Additional details are available in Table tablecountryperf

| country to predict | observed category | supersensitive | sensitive | resilient | rate | error |
|---|---|---|---|---|---|---|
| Brasil Mexico | supersensitive | **16** | 12 | 24 | 36 / 52 | 0.69 |
| Brasil Mexico | sensitive | 40 | **12** | 51 | 91 / 103 | 0.88 |
| Brasil Mexico | resilient | 16 | 9 | **44** | 25 / 69 | 0.36 |
| **Brasil Mexico** | **Totals** | **72** | **33** | **119** | **152 / 224** | **0.68** |
| Ghana Uganda | supersensitive | **100** | 23 | 10 | 33 / 133 | 0.25 |
| Ghana Uganda | sensitive | 60 | **21** | 21 | 81 / 102 | 0.79 |
| Ghana Uganda | resilient | 49 | 13 | **126** | 62 / 188 | 0.33 |
| **Ghana Uganda** | **Totals** | **209** | **57** | **157** | **176 / 423** | **0.42** |
| Kazakhstan India Poland | supersensitive | **20** | 35 | 29 | 64 / 84 | 0.76 |
| Kazakhstan India Poland | sensitive | 8 | **39** | 58 | 66 / 105 | 0.63 |
| Kazakhstan India Poland | resilient | 7 | 24 | **109** | 31 / 140 | 0.22 |
| **Kazakhstan India Poland** | **Totals** | **35** | **98** | **196** | **161 / 329** | **0.49** |
| England | supersensitive | **5** | 8 | 3 | 11 / 16 | 0.69 |
| England | sensitive | 36 | **74** | 14 | 50 / 124 | 0.40 |
| England | resilient | 29 | 76 | **81** | 105 / 186 | 0.56 |
| **England** | **Totals** | **70** | **158** | **98** | **166 / 326** | **0.51** |
| Uganda | supersensitive | **50** | 24 | 18 | 42 / 92 | 0.46 |
| Uganda | sensitive | 11 | **15** | 23 | 34 / 49 | 0.69 |

Table S6: Confusion matrices associated with the multinomial classification model (three categories) splitting the dataset into train and validation sets by country combinations. In the last row, we report the average error rate of 0.49. Additional details are available in Table tablecountryperf

| **country to predict** | **observed category** | **supersensitive** | **sensitive** | **resilient** | **rate** | **error** |
|---|---|---|---|---|---|---|
| Uganda | resilient | 10 | 17 | **88** | 27 / 115 | 0.23 |
| **Uganda** | **Totals** | **71** | **56** | **129** | **103 / 256** | **0.40** |
| **overall** | | | | | | **0.50** |

### Table S7: Comparison with forest-dependent species

Table S7: Percentage of species independently classed as having medium or high forest dependency ('forest dependent species') by density-yield response category. In the observed dataset, we include each species-by-study observation separately. In the modelled dataset, we assign the category using the highest probability value between the three estimated probabilities.

| | Supersensi | Sensitive | Resilient |
|---|---|---|---|
| **Observed (n = 1,008) % forest dependent species** | 40.7 | 38.5 | 20.8 |
| **Modelled (n = 9,530) % forest dependent species** | 70.3 | 20.4 | 9.3 |
| **Combined (n = 10,538) % forest dependent species** | 67.5 | 22.1 | 10.4 |

### Table S8: Mathematical symbols

Table S8: Mathematical symbols used in the equations and their definitions

| symbol | description |
|---|---|
| $t_i$ | scenarios $t_1$ observed values in 2015, $t_2$ projections in 2050 under business as usual, and $t_3$ projections in 2050 under closing yield gaps |
| $t_0$ | pristine habitat corresponding to 0 yields |
| $k$ | category $k_1$ (supersensitive), $k_2$ (Sensitive), or $k_3$ (resilient) |
| $s$ | a species |
| $p_{k_{1,2,3}}$ | probabilities of $s$ to belong to category $k_1$, $k_2$ and $k_3$. $p_{k_1} + p_{k_2} + p_{k_3} = 1$ |
| $c$ | a cell |
| $x_{c,t_i}$ | yield in cell $c$. $x_{c,t_0} = 0$ natural habitat; $x_{c,t_i}$ agriculture under scenario $t_i$ |
| $\zeta_c$ | fractional size of cell $c$ ($0.1 < \zeta_c <= 1$) |
| $\Omega_{s,c,x_{c,t_i}}$ | predicted density of species $s$ in cell $c$ under agriculture scenario $t_i$ |
| $\Omega_{s,c,x_{c,t_0}}$ | predicted density of species $s$ in cell $c$ in pristine habitat |
| $\omega_{k_{1,2,3};x_{c,t_i}}$ | expected density of category $k_1$, $k_2$, and $k_3$ at yield $x_{c,t_i/t_0}$ |
| $\gamma_{s,c,t_i}$ | suitable fraction in the AOH for species $s$ in cell $c$ |
| $\Gamma_s$ | size of the AOH maps of species $s$ standardized between 1 and 100 |
| $\phi_{c,t_i}$ | binary value indicating if the cell $c$ under scenario $t_i$ is covered by crops (1) or not (0) |
| $\theta_{s,c,t_i}$ | impact of species $s$ in cell $c$ under scenario $t_i$ |
| $\Theta_{c,t_i}$ | sum of impacts of all $n$ species in cell $c$ under scenario $t_i$ |
| $\Xi_{c,t_i}$ | proportion of species with overall density decline in cell $c$ under scenario $t_i$ |

### Table S9: Classification of species observed in multiple sites

Table S9: Pair-wise comparison of the classification of the 209 species observed up to nine times in multiple sites

| category | supersensitive | sensitive | resilient |
|---|---|---|---|
| **supersensitive** | **12** | 61 | 41 |
| **sensitive** | - | **173** | 285 |
| **resilient** | - | - | 345 |

### Appendix S1: Estimating density-yield curves

Density-yield curves are equations linking densities of each bird species ($i$) to yield values in each study site ($j$) of two possible forms (Green et al. 2005; Phalan et al. 2011). Model A is defined by the following equation:

$$y_{i,j} = \exp\left[b_{0,i,j} + b_{1,i,j} x_j^{\alpha_{i,j}}\right]$$

while model B is defined as:

$$y_{i,j} = \exp\left[b_{0,i,j} + b_{1,i,j} x_j^{\alpha_{i,j}} + b_{2,i,j} x_j^{2\alpha_{i,j}}\right]$$

where $y_{i,j}$ is the predicted density of species $i$ in study site $j$, $x_j$ is the yield in study site $j$, $b_{0,i,j}$, $b_{1,i,j}$, $b_{2,i,j}$ and $\alpha_{i,j}$ are parameters.

To estimate the parameters' values, the data were analyzed using constrained maximum likelihood estimation with linear and non-linear equality constraints using either the Nelder-Mead or BFGS methods (function constrOptim.nl in the R package alabama, R Core Team 2024; Varadhan and Varadhan 2015). The parameter $\alpha$ was constrained between 0 and 4.6 (following Green et al. 2005; Phalan et al. 2011). All parameters were constrained such that the value of $y_{i,j}$ was always lower than 1.5× the maximum observed density in study site $j$ (see Phalan et al. 2011 for further details). Many combinations of initial parameter values were used to ensure the parameter estimates did not converge to a local minimum. As a first step, the results were ranked using the $-2\times$ log-likelihood from lowest to largest values within model A (equation A) and model B (equation B), separately. When both model A and model B produced converged estimates, the results of models A and B were compared, and model B was preferred if the difference in the $-2\times$ log-likelihood between model A and model B was 3.84 or greater. Otherwise, model A was selected for reasons of parsimony.

Species that were only observed in natural habitats were assigned a density equal to the mean observed density in the natural habitat at zero yield, and a predicted density of zero for all other yields ($y_{i,j} = 0$ for any $x_j \geq 1$) (Phalan et al. 2011).

The study sites in the British Fens and Salisbury Plain include multiple types of natural vegetation (Finch et al. 2019). In the Fens, we assumed a composition of 1:1 of wet grassland and fen in pristine habitats, while in Salisbury Plain, a composition 1:1 of woodland and chalk grassland (Finch et al. 2019). As such, we included the density values in each natural habitat within each study site as single data points with equal weights. This assumption does not impact the overall results of the analysis, as shown in Figure S8 in Finch et al. (2019).

Feniuk, Balmford, and Green (2019) analyzed data collected in the Polesian lowlands in the Lubelskie region of eastern Poland. The natural vegetation in this environment is characterized by a mixture of forest (covering 75% of the site) and wetland (covering 25% of the site). The original study reported predicted bird density-yield curves in the two types of environment separately. Here, we combined the model predictions into a single density-yield curve by calculating the weighted mean between the two curves assuming that the environment was comprised of 75% forest and 25% wetland.

#### Additional details about the classification of species into categories

Supersensitive species were observed only in natural habitats or in agricultural lands with yields lower than 10% of the maximum yield observed in each study site or below the production target for the study site (whichever value is smaller). At all yields equal to or above this minimum threshold, the observed density of supersensitive species was zero. For species occurring only in natural habitats, only an intercept parameter could be estimated. The values for the slopes and exponent parameters were reported as missing.

Losers that fare better with land sparing (Sensitive) were found mainly in natural habitats and agriculture at low yields. As yields increase, the density of Sensitive species usually decreases steeply (convex curves). Sensitive species are defined as those which are not supersensitives but for which it is also the case that farming at maximum yield on the smallest possible area and protecting as much natural habitat as possible would result in the largest population size (Green et al. 2005).

We define as resilient all the species that have some tolerance for agriculture. The majority of these species have lower predicted density values in unfarmed habitat than under agriculture. Under any agricultural scenario, these species would have populations equal to, or larger than what their population would be in the absence of agriculture. We also include in the resilient categories, species that occur in both natural habitats and agriculture, often with their highest densities at intermediate yields. Their density typically declines at higher yields (88 species, concave curves, classified as land-sharing losers in other studies) (Green et al. 2005). These species are defined as those for which farming at the lowest permissible yield on the largest area possible would result in the largest population size. Finally, we include in the resilient category also the small number of species that occur in natural habitats but have their highest density at yields above the production target in the study site (7 species, classified as intermediate losers in other studies, Green et al. (2005)).

### Appendix S2: Formulaic definition of calculation of cell-level metrics of impact

We considered three possible time and agriculture management scenarios $t_i$: $t_1$ observed values in 2015, $t_2$ projected values in 2050 under business as usual, and $t_3$ projected values in 2050 with closing yield gaps (Williams et al. 2021). Between these three scenarios for each cell $c$, there could be changes in the yield values ($x_{c,t_i}$) or in the geographical distribution of crops ($\phi_{c,t_i}$) (variables are defined in Table S8). The impact $\theta_{s,c,t_i}$ of species $s$ in a cell $c$ under the three scenarios $t_i$ is defined as:

$$\theta_{s,c,t_i} = max[0, (\Omega_{s,c,x_{c,t_0}} - \Omega_{s,c,x_{c,t_i}})] \times (1 + \frac{\gamma_{s,c}\zeta_c}{\Gamma_s}) \times \phi_{c,t_i}$$

where the subscript $t_0$ indicates an hypothetical historical scenario with no agriculture; $\Omega_{s,c,x_{c,t_i}}$ and $\Omega_{s,c,x_{c,t_0}}$ are the predicted density of $s$ in cell $c$ at yield $x_{c,t_i} > 0$ and $x_{c,t_0} = 0$ respectively; $\gamma_{s,c}$ is the suitable fraction of a cell $c$ according to the AOH maps of species $s$; $\zeta_c$ is the fractional size of the cell ($\zeta_c = 1$ when the cell is land, while $0.1 < \zeta_c < 1$ in coastal areas); $\Gamma_s$ is the standardized size of the AOH maps of species $s$; and $\phi_{c,t_i}$ measure the percentage of the cell $c$ under scenario $t_i$ covered by crops (Table S8). The values of $\phi_{c,t_i}$ are re-projected to the grid 5x5 km from the results of Williams et al. (2021), who mapped the percentage of land used by crops at 2.25 km$^2$. In a migratory species, $\Gamma_s$ is set to the minimum between the size of the breeding and non-breeding AOH maps. The values of $\Gamma_s$ are standardized between 1 (the smallest AOH observed) and 100 (the largest AOH observed). The difference between the density in pristine habitat versus cropland, $\Omega_{s,c,x_{c,t_0}} - \Omega_{s,c,x_{c,t_i}}$ is positive for species that decline in density under cropland, negative for species that increase in density under agriculture, and zero when there is no change in density. To give lower weights to species that increases in density under agriculture, we truncate negative values of impact ($\theta_{s,c,t_i}$) to 0.

According to the observations or the predictions from the classification model, each species is assigned three probabilities of belonging to each strategy: $p_{k=1,2,3}$. The sum of the three probabilities is 1, $\sum_{k=1}^{3} p_k = 1$.

The predicted density $\Omega_{s,c,x_{c,t_i}}$ of species $s$ in cell $c$ is the expected density of the three categories $\omega_{k_{1,2,3},x_{c,t_i}>0}$ at yield $x_{c,t_i}$ weighted by the species-specific probabilities $p_{k_{1,2,3}}$ to belong to the three categories of $s$ and equals:

$$\Omega_{s,c,x_{c,t_i}} = p_1\omega_{k_1,x_{c,t_i}} + p_2\omega_{k_2,x_{c,t_i}} + p_3\omega_{k_3,x_{c,t_i}}$$

The predicted density $\Omega_{s,c,x_{c,t_0}=0}$ of species $s$ in cell $c$ if the cell is covered by pristine habitats ($x_{c,t_0}$) equals:

$$\Omega_{s,c,x_{c,t_0}} = p_1 + p_2 + p_3 0.073$$

because the predicted densities for supersensitive and sensitive species at 0 yields (i.e., $t_0$) is $\omega_{k_{1,2},x_{c,t_0}} = 1$ and for resilient is $\omega_{k_3,x_{c,t_0}} = 0.073$. Therefore, the value of $\Omega_{s,c,x_{c,t_0}}$ ranges between 0.073 (when a species is a resilient with $p_{k_3} = 1$) and 1 (when ($p_{k_1} + p_{k_2} = 1$).

We summed all the impact values, $\theta_{s,c,t_i}$ of all species occupying each cell to calculate the overall impact of each cell, which equals:

$$\Theta_{c,t_i} = \sum_{s=1}^{n} \theta_{s,c,t_i}$$

where $n$ is the total number of species in the cell $c$.

The mean impact in cell $c$ equals:

$$\mu(\Theta_{c,t_i}) = \frac{\Theta_{c,t_i}}{n}$$

and its standard deviation is:

$$\sigma(\Theta_{c,t_i}) = \sqrt{\frac{\sum(\Theta_{c,t_i} - \mu(\Theta_c, t_i))^2}{n}}$$

261 . The proportion of species with overall density declines in the cell $c$ equals

$$\Xi_{c,t_i} = \frac{\sum_{s=1}^{n} [(\text{if } \theta_{s,c,t_i} > 0 \rightarrow 1) \wedge (\text{if } \theta_{s,c,t_i} = 0 \rightarrow 0)]}{n}$$

### Appendix S3: Concordance between multiple assessments of density-yield category of the same species

Of the 862 species recorded across all study sites, 209 were recorded at two or more sites (max = 9 sites) and hence had two or more independent density-yield category assessments. Between them, these 209 species yielded a total of 917 pairwise category assessment comparisons (e.g. a species recorded at 4 sites (a, b, c and d) would yield 6 pairwise comparisons between sites: a vs b, a vs c, a vs d, b vs c, b vs d, and c vs d). Across these 917 species-by-site pairwise comparisons, 530 (57.8%) were consistent between sites (S9).

To assess the concordance in classification between sites, we calculated the index of inter-rater reliability for ordinal ratings (Kendall's coefficient of concordance W) for unbalanced datasets (Kendall 1962; Taylor 1987). The index of interrater reliability is traditionally used to measure the concordance between the classification of an entity to a class by multiple raters with unbalanced datasets (i.e., some entities were not assessed by all raters). To calculate the coefficient, we converted the three impact categories into an ordinal variable (supersensitive = 1, Sensitive = 2, and resilient = 3), and we considered each site as a different rater. Kendall's coefficient of concordance, W, is a weighted mean of all Spearman's pair-wise comparisons that varies between 0 and 1 (Kendall 1962; Taylor 1987) and is interpreted subjectively. We obtained a value of 0.54, which is generally taken to indicate 'moderate agreement' between sites (Landis and Koch 1977)

### Appendix S4: Validation of density-yield response categories using the Human Tolerance Index (HTI) of Marjakangas et al. 2024

The Human Tolerance Index or HTI (Marjakangas et al. 2024) is derived at the continental scale from eBird occurrences as a function of the human footprint index. The human footprint index includes agriculture along with other pressures, such as urbanization. The conservative HTI represents the species' upper limits of tolerance to human disturbance. The peak HTI is the optimal human footprint index for the species. Further details are available in Marjakangas et al. (2024). The authors showed that 22% of birds can tolerate high human modifications (human footprint index $> 40$), while 0.001 percent only tolerate intact environments (human footprint index $< 4$). We compared our three density-yield response categories with values of the conservative (Figure S6 A1 and A2) and peak (Figure S6 B1 and B2) HTI. In the first row, we show the comparison using only the observed dataset, which includes information about the continent where the species was observed (to match the HTI's continental specificity) (Figure S6 A1 and B1). In the second row, we compare all species (i.e., those whose density-yield response category was observed and those for which it was imputed by our deep-learning models) with the HTI metrics ignoring the continent (Figure S6 A2 and B2). On the y-axes we plot the density of each category of impact to agriculture against the two HTI metrics. In the observed dataset, we could assign the continent to each observation and each species-by-site classification was analysed separately. In the predicted dataset, we could not discriminate the continent and we assigned the category using the highest probability values between the three probabilities of belonging to each category. The majority of supersensitive species have lower than average HTI and resilient species have higher HTI. Sensitive species show density distributions somewhat intermediate between supersensitive and resilient species, but close to those of resilient species. This could be because the HTI considers multiple human impacts and not solely agriculture; it seems plausible that Sensitive species have higher tolerance to other forms of disturbance than do supersensitive species.

### Appendix S5: Software details

The results were obtained using R Core Team (2024) and GDAL/OGR (2024). The list of R packages loaded can be found below.

```
# list of used libraries
if (!require("pacman")) install.packages("pacman")
pacman::p_load(foreach, doSNOW, leaflet,
  sf, ggplot2, sp,
  rnaturalearth, rnaturalearthdata,
  tidyverse, data.table, vioplot,
  viridis, foreign, terra, fst, collapse,
  Rfast, R.utils, Rcpp, hrbrthemes,
  terra, janitor, dplyr, stringr,
  gdalUtilities, collapse,
  ggplot2, plyr, h2o, bigvis)
```

All scripts needed to reproduce the analysis are archived XXX

Marjakangas, Emma-Liina, Alison Johnston, Andrea Santangeli, and Aleksi Lehikoinen. 2024. "Bird Species' Tolerance to Human Pressures and Associations with Population Change." *Global Ecology and Biogeography*.

Phalan, Ben, Malvika Onial, Andrew Balmford, and Rhys E Green. 2011. "Reconciling Food Production and Biodiversity Conservation: Land Sharing and Land Sparing Compared." *Science* 333 (6047): 1289–91.

R Core Team. 2024. "R: A Language and Environment for Statistical Computing." Vienna, Austria: R Foundation for Statistical Computing. https://www.R-project.org/.

Taylor, Jeremy MG. 1987. "Kendall's and Spearman's Correlation Coefficients in the Presence of a Blocking Variable." *Biometrics*, 409–16.

Varadhan, Ravi, and Maintainer Ravi Varadhan. 2015. "Package 'Alabama'."

Williams, David R, Michael Clark, Graeme M Buchanan, G Francesco Ficetola, Carlo Rondinini, and David Tilman. 2021. "Proactive Conservation to Prevent Habitat Losses to Agricultural Expansion." *Nature Sustainability* 4 (4): 314–22.
